# Fibronectin silencing remodels the fibrotic bone marrow niche

**DOI:** 10.64898/2026.10.16.759614

**Authors:** Francesca Rossella Calledda, Vittorio Abbonante, Maria Francesca Testa, Delfina Tosi, Carla Martinelli, Santo Diprima, Roberta Gioia, Umberto Gianelli, Caroline Marty, Hana Raslova, Isabelle Plo, Dario Balestra, Mirko Pinotti, Alessandra Balduini, Alessandro Malara

**Author notes:** To whom correspondence should be addressed: Alessandro Malara, Department of Molecular Medicine, University of Pavia, Campus della Salute, Viale Golgi 19, 27100, Pavia, Italy.

## Abstract

Bone marrow (BM) fibrosis is a defining feature of Primary Myelofibrosis, yet the extracellular matrix (ECM) that sustains the fibrotic niche remains largely untargeted therapeutically. Fibronectin (FN), a key organizer of ECM architecture and growth factor signaling, accumulates in the fibrotic BM niche and correlates with disease severity. Here, we investigated whether systemic FN depletion can disrupt pathological remodeling of the BM niche. We developed a chemically modified antisense oligonucleotide targeting *FN1* (FN ASO), which reduced FN expression and assembly and impaired transforming growth factor-β1 (TGF-β1)-induced activation of mesenchymal stromal cells. Systemic FN ASO administration efficiently depleted circulating and BM FN in mice and was well tolerated during short-term treatment. In two complementary models of BM fibrosis, FN silencing reduced FN matrix organization, stromal activation, active TGF-β1, reticulin, and collagen deposition. In *Jak2*^V617F^-driven disease, transcriptomic profiling further revealed attenuation of ECM, TGF-β1, and mesenchymal remodeling programs, accompanied by changes in inflammatory signaling. Collectively, these findings identify FN as an active organizer of the fibrotic BM niche and show that its systemic silencing uncouple ECM organization from profibrotic stromal signaling, providing rationale for targeting FN in myelofibrosis with broader therapeutic relevance to fibrosis-driven diseases.

## Introduction

Bone marrow fibrosis (MF) is a major pathological feature of primary myelofibrosis (PMF), a Philadelphia chromosome-negative myeloproliferative neoplasm characterized by clonal hematopoiesis, progressive remodeling of the bone marrow (BM) microenvironment, splenomegaly, and ineffective hematopoiesis^1,2^. Activating mutations in *JAK2, CALR*, or *MPL* drive constitutive JAK-STAT signaling in the malignant clone^3,4^, while abnormal megakaryocytes and inflammatory mediators promote stromal activation and excessive extracellular matrix (ECM) deposition^5^. JAK inhibitors improve splenomegaly and disease-related symptoms, but their impact on BM fibrosis remains limited^6^, highlighting the need for strategies that directly target the pathological niche.

Fibronectin (FN), encoded by the *FN1* gene, is a central regulator of ECM organization and cell fate, controlling adhesion, migration, and differentiation through integrin-dependent signaling. FN exists as a soluble plasma form, primarily produced by hepatocytes, and as a tissue associated form that is assembled into insoluble ECM fibrils^7^. Although FN expression is limited in most healthy adult tissues, it is rapidly induced following tissue injury and in fibroproliferative disorders. Importantly, FN contributes to the regulation of transforming growth factor-β1 (TGF-β1) signaling by facilitating the incorporation and organization of latent TGF-β complexes within the ECM, thereby influencing TGF-β1 bioavailability, activation, and downstream receptor signaling^8,9^. In fibrotic conditions, excessive FN deposition and altered matrix organization can therefore sustain TGF-β1-dependent myofibroblast activation and pathological ECM remodeling, contributing to the establishment of a self-reinforcing profibrotic microenvironment^10–12^. Circulating FN can also be incorporated into tissue matrices, indicating that both systemic and locally produced FN may contribute to fibrotic remodeling^13,14^.

FN accumulates in PMF, but whether FN is required to maintain the profibrotic niche, and whether depletion of both circulating and locally produced FN can modify fibrosis, is unknown. Aberrant FN expression and secretion have been reported in mesenchymal stromal cells (MSCs) from patients with PMF^15^, and BM FN levels correlate with fibrosis grade and disease severity^16^. In particular, the alternatively spliced EDA-FN isoform is detected during fibrosis progression in both plasma and BM biopsies from PMF patients and in murine models of MF^16,17^. Moreover, targeting the α5 subunit of the α5β1 integrin, a major FN receptor, has been proposed as a therapeutic strategy to limit disease progression in Jak2^V617F⁺^ mice^18^. Here, we tested whether systemic FN silencing could remodel the fibrotic BM niche using an *FN1*-directed antisense oligonucleotide in cellular systems and complementary mouse models of MF.

## Results

### FN silencing disrupts stromal activation and extracellular matrix assembly

We designed a gapmer with 2’-O-methyl (2′OMe) wings and phosphorothioate (PS) backbone antisense oligonucleotide (ASO) that simultaneously suppresses the expression of both cellular and plasma FN forms **(Figure 1A)**. Dose-response studies showed that FN ASO significantly reduced *FN1* mRNA levels in primary human MSCs **(Figure 1B)** and hepatic cell lines (Huh-7 and Hepa 1-6) **(Figure S1A-B)** compared with scramble ASO, with significant suppression at 150 nM, which was selected for subsequent *in vitro* experiments. FN ASO effectively targeted *EDA-* and *IIICS*-containing transcripts as evaluated by PCR splicing assay in both untreated and TGF-β1-treated mouse embryonic fibroblasts **(Figure 1C)**.

**Figure 1.**
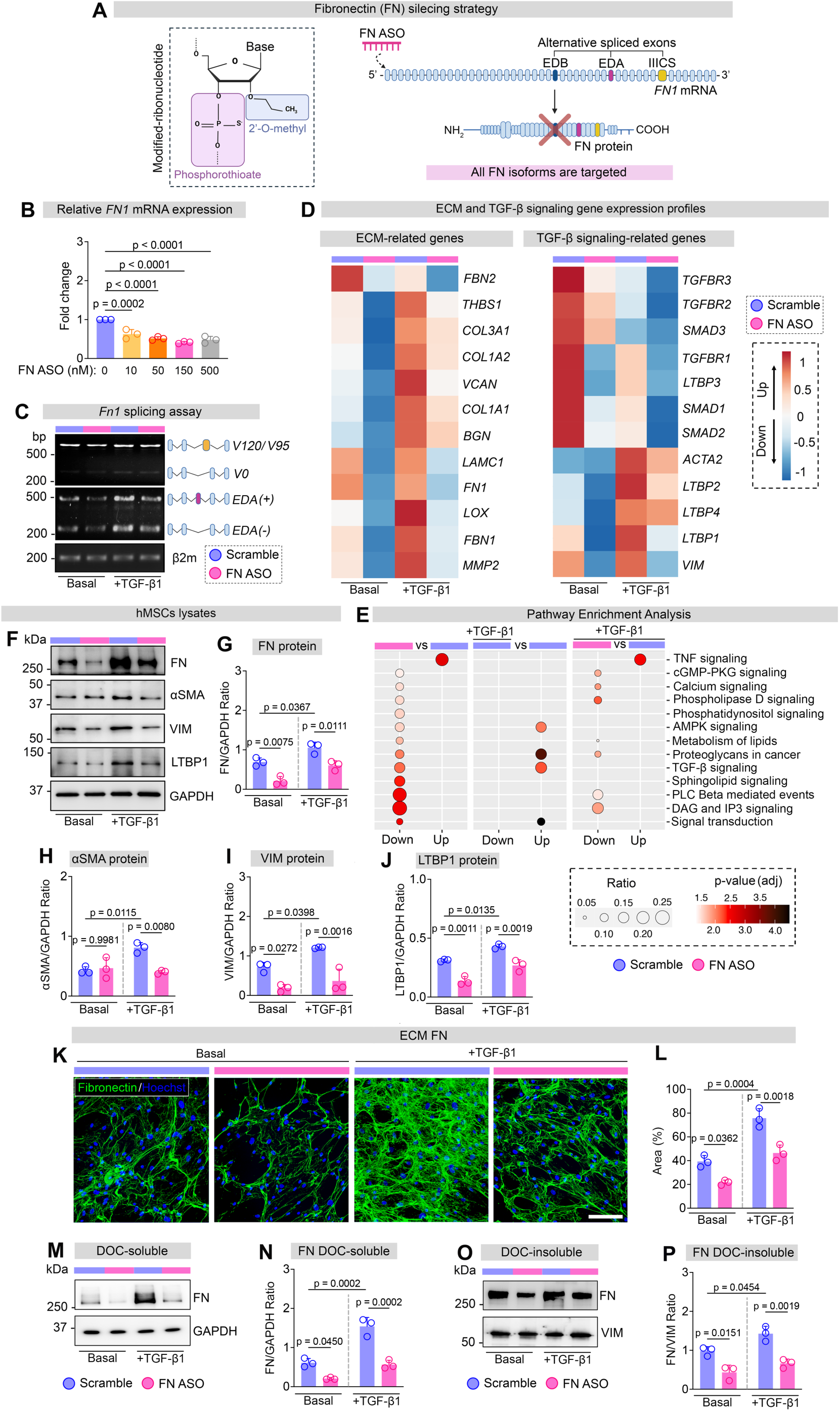
Fibronectin is required for fibrotic activation and matrix assembly in mesenchymal stromal cells. **A)** Schematic illustration of the chemical modifications of FN ASO and its mechanism of action for silencing FN expression. **B)** Dose-dependent effects of FN ASO on *Fn1* mRNA expression in hMSCs (n = 3). Gene expression was normalized to GAPDH and expressed relative to the untreated group, which was set to 1. Ordinary one-way ANOVA was used. **C)** PCR-splicing assay of *EDA-* and *IIICS*-containing FN isoforms in murine embryonic fibroblasts (mEFs) transfected with FN ASO or control scrambled ASO, either untreated or stimulated with TGF-β1. **D)** Heatmap showing the average expression of ECM- and TGF-β1-related genes in hMSCs treated with scrambled ASO, FN ASO, TGF-β1 plus scrambled ASO, or TGF-β1 plus FN ASO (n = 4). Color intensity represents scaled gene expression (z-score). **E)** Pathway enrichment analysis of hMSCs treated with FN ASO versus scrambled ASO, scrambled ASO plus TGF-β1 versus scrambled ASO, and FN ASO plus TGF-β1 versus scrambled ASO plus TGF-β1 (n = 4). The ratio indicates the magnitude of pathway enrichment, while adjusted P values indicate statistical significance. **F-J)** Representative Western blot analysis of FN, α-SMA, Vimentin, and LTBP1 in whole-cell lysates from hMSCs treated with scrambled ASO, FN ASO, TGF-β1 plus scrambled ASO, or TGF-β1 plus FN ASO (F), with corresponding relative quantification (G-J) (n = 3). GAPDH served as the loading control. Ordinary two-way ANOVA was used. **K-L)** Immunofluorescence staining of FN (green) in hMSCs treated with scrambled ASO, FN ASO, TGF-β1 plus scrambled ASO, or TGF-β1 plus FN ASO (K), with corresponding quantification (L) (n = 3). Nuclei (blue) were counterstained with Hoechst 33258. Images were acquired using a 20x objective; scale bar = 50 μm. Ordinary two-way ANOVA was used. **M-P)** Western blot analysis of FN in DOC-soluble (M-N) and DOC-insoluble (O-P) fractions from hMSCs treated with scrambled ASO, FN ASO, TGF-β1 plus scrambled ASO, or TGF-β1 plus FN ASO, with corresponding relative quantification (N and P) (n = 3). GAPDH and vimentin were used as loading controls for the DOC-soluble and DOC-insoluble fractions, respectively. Ordinary two-way ANOVA was used.

As PS-modified ASOs can trigger innate immune responses through Toll-like receptor 9 (TLR9), we next assessed whether FN ASO could activate TLR9 signaling. FN ASO did not induce detectable TLR9 activation in HEK-Blue^TM^ reporter cells expressing mouse TLR9, whereas ODN1668, a well-established B-class CpG oligodeoxynucleotide agonist of mouse TLR9, elicited a robust response **(Figure S1C**). FN depletion also did not cause apoptosis or measurable cell death in viability assay **(Figure S1D)**. To assess the impact of FN targeting on the profibrotic phenotype of fibrosis-driving cells, bulk RNA sequencing was performed on MSCs treated with FN ASO or scramble control under basal conditions or following stimulation with TGF-β1 for 24 hours. In addition to *FN1* gene, FN depletion broadly suppressed transcriptional programs associated with ECM organization and stromal activation, including ECM components (e.g. *FBN1, THBS1, VCAN, BGN, COL1A2, COL3A1*), ECM remodeling factors (e.g. *LOX, MMP2*), and mesenchymal-to-myofibroblast transition markers and signaling (e.g. *ACTA2, VIM, TGFBR1, TGFBR2, LTBP1*) **(Figure 1D and Table S1)**. Gene ontology and pathway enrichment analyses were performed to investigate the transcriptional programs modulated by FN silencing. Silencing of *FN1* with ASO revealed a marked suppression of signaling pathways associated with profibrotic activation, including TGF-β1 signaling, proteoglycans in cancer, and multiple pathways involved in intracellular signal transduction such as phosphatidylinositol signaling, PLCβ-mediated DAG/IP3 signaling, and calcium signaling, suggesting that FN contributes to the regulation of signaling networks controlling mesenchymal cell activation **(Figure 1E and Table S2)**. Importantly, the attenuation of some of these signaling programs persisted in TGF-β1-stimulated MSCs, indicating that the ASO retains its ability to disrupt cellular activation even under profibrotic conditions **(Figure 1E)**. Upregulated signaling was primarily enriched for pathways associated with cellular splicing and TNF signaling, indicating that FN depletion does not uniformly suppress inflammatory signaling. The ability of FN ASO to reduce FN protein deposition and the acquisition of a TGF-β1-dependent activated profibrotic phenotype (e.g. α-Smooth Muscle Actin (α-SMA), Latent TGF-β binding protein-1 (LTBP1) and Vimentin (Vim) levels) was further validated by western blot and immunofluorescence analyses of treated-cells, respectively **(Figure 1F-J and Figure 1K-L)**. FN assembly was also significantly impaired, as indicated by reduced incorporation of FN in the cell-associated deoxycholic acid (DOC)-soluble fraction **(Figure 1M-N)** and in the DOC-insoluble matrix fraction **(Figure 1O-P)**.

Moreover, stromal cells treated with FN ASO showed reduced cell migration **(Figure S1E-F)** and contractility **(Figure S1G-H)** in a wound healing and collagen gel contraction assay, respectively, even upon TGF-β1 stimulation. Overall, these data reveal that lowering FN expression and deposition mitigates profibrotic signaling, ECM accumulation, and myofibroblast activation, highlighting FN as a key regulator of stromal remodeling.

### Systemic FN ASO efficiently depletes circulating and tissue FN in vivo

To assess the targeting efficacy of FN ASO and its safety profile *in vivo*, steady-state wild-type C57BL/6 mice were injected five times during two weeks with FN ASO at 50 mg/kg and sacrificed at day 14 (**Figure 2A**). The dose was selected based on previous studies on therapeutic ASOs^19^. Plasma FN levels were significantly reduced, whereas levels of another liver-derived plasma protein, transferrin, remained unchanged **(Figure 2B-E)**. Immunostaining of BM and liver sections using an anti-PS antibody revealed accumulation of FN ASO within these tissues at sacrifice, suggesting sustained target-tissue uptake **(Figure 2F)**. No mortality or body weight changes **(Figure S2A)** were observed and most blood parameters were unchanged after 2 weeks, except for a modest but significant reduction in platelet counts **(Figure S2B)**. Plasma fibrinogen levels **(Figure S2C)**, coagulation assays (Prothrombin (PT) and Activated Partial Thromboplastin Time (APTT)) **(Figure S2D)**, agonist-dependent platelet activation (ADP and Collagen) **(Figure S2E)** and blood markers of tissue damage (alanine aminotransferase (ALT-GPT), aspartate aminotransferase (AST-GOT), albumin, total bilirubin, lactate dehydrogenase (LDH), urea) remained within normal ranges **(Figure S2F)**. Lower *Fn1* mRNA expression in BM and liver tissues was confirmed by qPCR **(Figure S2G)**. Unspliced and alternatively-spliced *Fn1* transcripts were similarly reduced in liver and BM samples of FN-depleted mice **(Figure S2H)**. Consistently, analysis at sacrifice showed a significant reduction in FN protein expression and deposition in both liver **(Figure 2G-J)** and BM **(Figure 2L-P)**, as assessed by immunoblotting and immunofluorescence, respectively. Transmission electron microscopy of liver and BM revealed increased intercellular spaces and supportive ultrastructural evidence of reduced reticular connective tissue exclusively in FN ASO-treated mice **(Figure S2I)**. Overall, these findings confirmed that FN ASO can systemically lower both pools of FN without major acute toxicity. BM immunophenotyping revealed only minor changes in the relative frequencies of hematopoietic lineages, as assessed by flow cytometry. Erythroid profiling showed a selective increase in basophilic erythroblasts (R2), while early and late stages were unchanged **(Figure S2J**). No major differences were observed in major myeloid and lymphoid lineages **(Figure S2J)**. Given the contribution of systemic FN to bone mass homeostasis^20^, we also assessed bone mass in treated mice. Micro-computed tomography (μCT) analysis of cortical and trabecular bone structure revealed no significant differences in bone volume, thickness, or mineral density **(Figure S2K)**. Together, these findings indicate that systemic FN suppression is sufficient to remodel the ECM *in vivo*, without overt biochemical or hematological evidence of acute organ toxicity over the treatment period.

**Figure 2.**
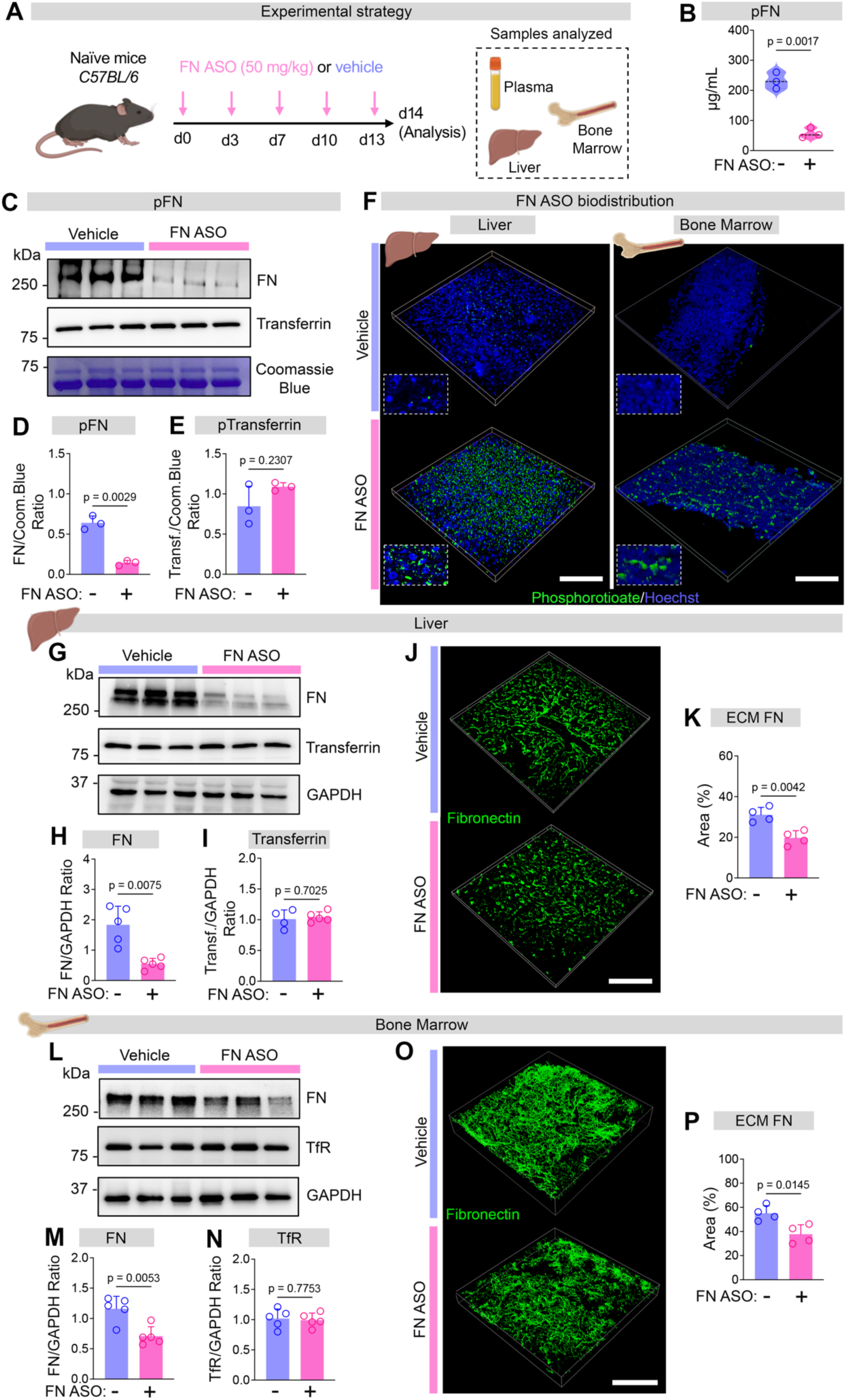
Systemic FN ASO efficiently depletes circulating and tissue-associated FN. **A)** Schematic representation of the experimental design for FN ASO or vehicle administration in mice. **B)** ELISA quantification of plasma FN (pFN) levels in mice treated with vehicle (n = 4) or FN-ASO (n = 3). Welch’s t test was used. **C-E)** Representative western blot analysis of FN and transferrin in plasma samples from vehicle- and FN ASO-treated mice (C), with corresponding densitometric quantification (D-E) (n = 3). Coomassie Brilliant Blue staining served as the loading control. Welch’s t test was used. **F)** Representative immunofluorescence images showing phosphorothioate (PS) staining (green) in liver and bone marrow (BM) sections from vehicle- and FN ASO-treated mice. Images were acquired using a 25x objective; scale bar = 200 μm. **G-I)** Representative western blot analysis of FN, transferrin, and transferrin receptor (TfR) in liver lysates from vehicle- and FN ASO-treated mice (G), with corresponding densitometric quantification (H-I) (n = 5). GAPDH served as the loading control. Welch’s t test was used. **J-K)** Immunofluorescence staining of FN in liver sections from vehicle- and FN ASO-treated mice (J), with corresponding quantification (K) (n = 4). Images were acquired using a 25x objective; scale bar = 200 μm. Each data point represents one biological replicate and corresponds to the mean percentage of positive area measured across at least five independent fields. Welch’s t test was used. **L-N)** Representative western blot analysis of FN, transferrin, and transferrin receptor (TfR) in BM lysates from vehicle- and FN ASO-treated mice (L), with corresponding densitometric quantification (N) (n = 5). GAPDH served as the loading control. Welch’s t test was used. **O-P)** Immunofluorescence staining of FN in BM sections from vehicle- and FN ASO-treated mice (O), with corresponding quantification (P) (n = 4). Images were acquired using a 25x objective; scale bar = 200 μm. Each data point represents one biological replicate and corresponds to the mean percentage of positive area measured across at least five independent fields. Welch’s t test was used.

### Systemic FN silencing limits the development of Romiplostim-induced bone marrow fibrosis

MF can be pharmacologically induced by Romiplostim injections. Romiplostim, a thrombopoietin receptor agonist used clinically to treat thrombocytopenia, when administered at supra-pharmacological doses, recapitulates key features of myelofibrosis in mice including splenomegaly, increased megakaryocyte expansion, and elevated levels of active TGF-β1 within the BM. Using this model, we confirmed that plasma FN contributes to FN accumulation during fibrosis progression. Indeed, intravenously injected fluorescently labelled human plasma FN accumulated and was incorporated into visible fibrils within the BM matrix exclusively under fibrotic conditions, whereas labelled albumin (negative control) was rapidly cleared out from the BM environment **(Figure 3A-C)**. These findings suggest that circulating FN is not merely elevated during fibrosis but is actively recruited and incorporated into the pathological BM matrix. To evaluate the therapeutic potential of FN ASO, mice received injections of FN ASO (50 mg/kg) or vehicle (PBS) for 14 days before and for 14 days during pharmacological induction of BM fibrosis with Romiplostim **(Figure 3D)**. FN ASO significantly reduced plasma FN levels in Romiplostim-treated mice **(Figure 3E),** without affecting body weight **(Figure S3A)**. Notably, active and total TGF-β1 level in BM cell-free fluids were reduced following FN ASO treatment, as determined by ELISA and immunoblot analyses **(Figure 3F-H)**.

**Figure 3.**
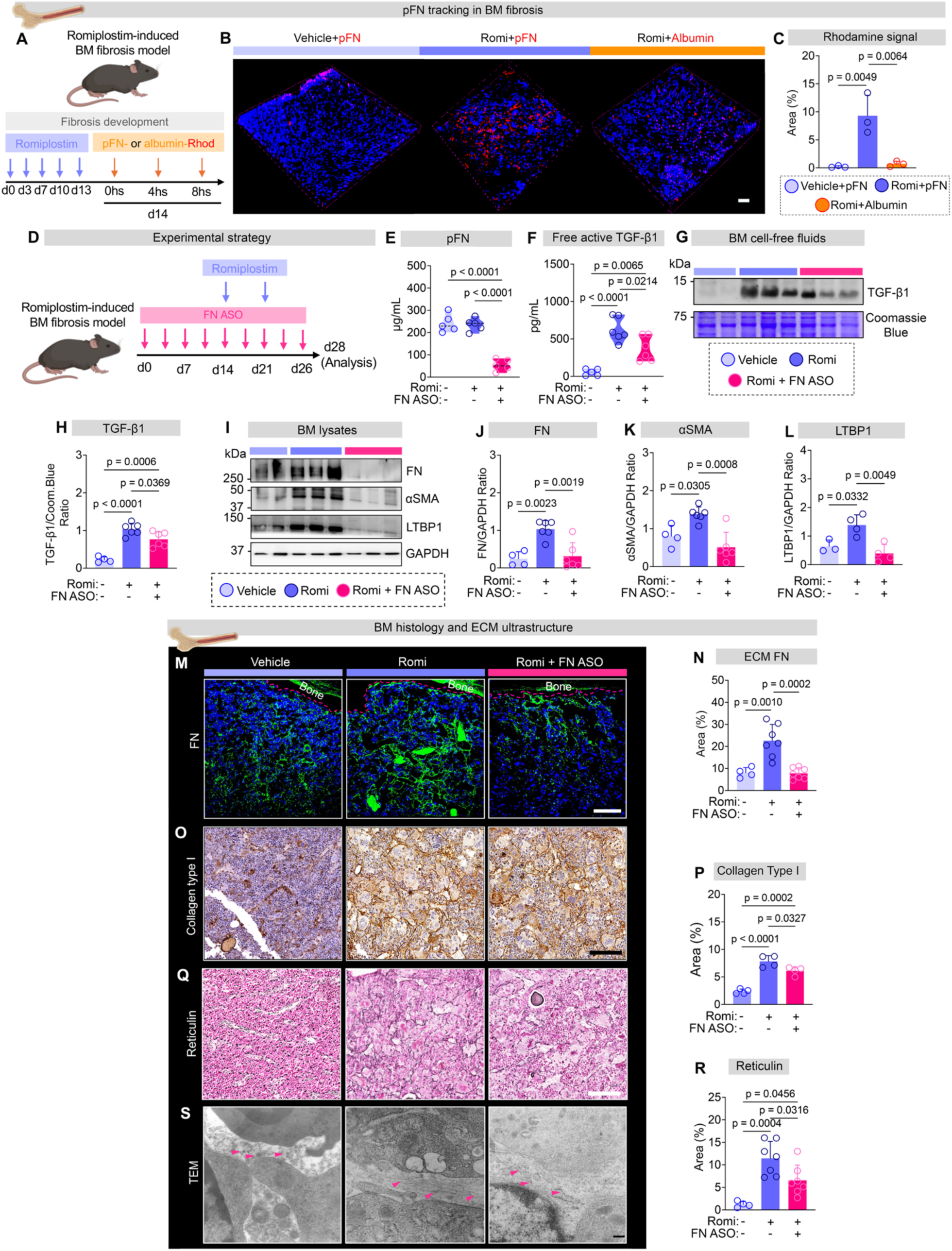
Circulating FN is incorporated into the fibrotic BM matrix and FN silencing suppresses Romiplostim-dependent fibrotic remodelling. **A-C)** Immunofluorescence analysis of BM sections following intraperitoneal administration of rhodamine-conjugated plasma FN or albumin (red signal) in vehicle- and Romiplostim-treated mice (A-B), with quantification of the area covered by the fluorescent signal (C) (n = 3). Images were acquired using a 25x objective; scale bar = 50 μm. Each data point represents one biological replicate and corresponds to the mean percentage of positive area measured across at least five independent fields. Ordinary two-way ANOVA was used. **D)** Schematic representation of the experimental design for BM fibrosis induction and FN ASO administration in mice. **E)** ELISA quantification of plasma FN levels in vehicle-treated (n = 5), Romiplostim plus vehicle-treated (n = 5), and Romiplostim plus FN-ASO-treated (n = 5) mice. Ordinary two-way ANOVA was used. **F)** ELISA quantification of active TGF-β1 levels in BM cell-free fluids from vehicle-treated (n = 5), Romiplostim plus vehicle-treated (n = 6), and Romiplostim plus FN-ASO-treated (n = 6) mice. Ordinary two-way ANOVA was used. **G-H)** Representative western blot analysis of TGF-β1 in BM cell-free fluids from vehicle-treated (n = 4), Romiplostim plus vehicle-treated (n = 6), and Romiplostim plus FN ASO-treated (n = 6) mice (G), with corresponding relative quantification (H). Coomassie Brilliant Blue staining served as the loading control. Ordinary two-way ANOVA was used. **I-L)** Representative western blot analysis of FN, LTBP1, and α-SMA in BM cell lysates from vehicle-, Romiplostim plus vehicle-, and Romiplostim plus FN ASO-treated mice (I), with corresponding relative quantification (J-L). At least four biological replicates were analyzed. GAPDH served as the loading control. Ordinary two-way ANOVA was used. **M-N)** Immunofluorescence staining of FN (green) in BM sections from vehicle-treated (n = 3), Romiplostim plus vehicle-treated (n = 7), and Romiplostim plus FN ASO-treated (n = 7) mice (M), with corresponding quantification (N). Nuclei (blue) were counterstained with Hoechst 33258. Images were acquired using a 25x objective; scale bar = 150 μm. Each data point represents one biological replicate and corresponds to the mean percentage of positive area measured across at least five independent fields. Ordinary two-way ANOVA was used. **O-P)** Representative immunohistochemical staining of type I collagen in BM sections from vehicle-Romiplostim+vehicle- and Romiplostim+FN ASO-treated mice (O) with quantification of the area covered by type I collagen (P) (n=4). Ordinary two-way ANOVA was used. **Q-R)** Reticulin staining of BM sections from vehicle-treated (n = 4), Romiplostim plus vehicle-treated (n = 7), and Romiplostim plus FN ASO-treated (n = 7) mice (Q), with corresponding quantification of the area covered by reticulin staining (R). Images were acquired using a 20x objective; scale bar = 100 μm. Each data point represents one biological replicate and corresponds to the mean percentage of positive area measured across at least five independent fields. Ordinary two-way ANOVA was used. **S)** Transmission electron microscopy images of BM from vehicle-, Romiplostim plus vehicle-, and Romiplostim plus FN ASO-treated mice. Pink arrows indicate ECM components. Scale bar = 500 nm.

Consistent with these findings, FN ASO reduced the expression of FN, α-SMA, and LTBP1 in the BM at the protein level, suggesting reduced stromal activation and attenuation of the TGF-β signaling axis **(Figure 3I-L).** Red blood cell and leukocyte counts, hematocrit, and hemoglobin levels remained unchanged **(Figure S3B)**. No differences were detected in the distinct stages of terminal erythroid differentiation **(Figure S3C)**, while, FN silencing attenuated Romiplostim-induced thrombocytosis and reduced BM megakaryocyte content and size **(Figure S3B, S3D-F).** Importantly, immunofluorescence and immunohistochemistry analyses further demonstrated reduced FN accumulation **(Figure 3M-N)**, type I collagen (**Figure 3O-P**), and reticulin deposition in the BM of FN ASO-treated mice (**Figure 3Q-R**). Transmission electron microscopy analysis confirmed an overall reduction and loosening of the BM fibrotic matrix following FN ASO treatment (**Figure 3S)**. Spleen/body weight ratio was unchanged at sacrifice **(Fig. S3G-H)**. Extramedullary hematopoiesis persisted in FN ASO-treated mice, although erythroid progenitor frequencies shifted toward those observed in control wild-type mice, particularly within the basophilic erythroblasts (R2) and orthochromatophilic erythroblasts (R4) fractions **(Fig. S3I)**. Notably, markers of stromal activation, including α-SMA, FN and type III collagen were reduced at both the mRNA and protein levels **(Figure S3J-L)**, accompanied by decreased reticulin deposition **(Fig. S3M)**. Collectively, these findings demonstrate that FN silencing exerts a marked anti-fibrotic effect in the Romiplostim-induced model of BM fibrosis, primarily by attenuating stromal activation, TGF-β1 availability, and ECM deposition, despite only partial correction of the associated hematopoietic abnormalities.

### FN silencing remodels the fibrotic niche in Jak2^V617F^-driven myelofibrosis

Given the slow and variable fibrosis development in standard *Jak2*^V617F⁺^ mouse models we took advantage of the recently characterized KOKI mice. These mice are knock-out (KO) for p19^Ink4d^, a CDK4/CDK6 inhibitor that regulates the endomitotic arrest of megakaryocytes, and knock-in (KI) for the *Jak2*^V617F^ mutation (named KOKI) and display an accelerated progression of BM fibrosis (less than 3 weeks upon *Jak2*^V617F^ induction). To test the efficacy of the FN ASO we used a model of primary transplanted mice, with BM cells of the KOKI mice being engrafted into lethally-irradiated wild type mice. The treatment with FN ASO (50 mg/kg, five injections every two weeks) and vehicle (PBS) was initiated after *Jak2*^V617F^ mutation induction by Tamoxifen (1 week) and maintained until sacrifice at week 5 **(Figure 4A)**. Plasma and hepatic FN levels were significantly reduced in FN ASO-treated KOKI mice, while transferrin levels were unperturbed (**Figure 4B and Figure S4A**). Active TGF-β1 levels in BM cell-free fluids were significantly reduced **(Figure 4C)**. To investigate the molecular consequences of *Fn1* silencing in the BM of KOKI mice, we performed RNA-seq analysis of total BM cell suspensions following red blood cell lysis from KOKI mice treated with FN ASO or vehicle. PCA analysis confirmed clear separation among samples **(Figure S4B)**, while differential expression analysis identified 581 significantly downregulated and 645 upregulated genes following *Fn1* silencing **(Figure S4C)**. Consistent with the pharmacological targeting strategy, *Fn1* mRNA levels were significantly reduced in the BM of FN ASO-treated mice **(Figure S4D)**. Among the most prominently downregulated transcripts were genes associated with the ECM and TGF-β-associated fibrotic response, including *Thbs1, Serpine1, Fn1, Lama5, Smad7, Tgif1 and Tgif2* **(Figure 4D and Table S3)**, indicating broad remodeling of the ECM/TGF-β-associated transcriptional program following *Fn1* silencing. In parallel, inflammatory cytokines and chemokines, including *Il1b, Il6 and Cxcl2*, were significantly downregulated, whereas *Tnf* and *interferon-γ*-associated responses were increased, suggesting that *Fn1* silencing does not induce a generalized suppression of immune signaling but rather promotes a qualitative shift in the inflammatory state **(Figure 4D)**. Gene set enrichment analysis confirmed these findings and further revealed a prominent impact of *Fn1* silencing on pathways related stromal activation and differentiation **(Figure 4E)**. Among the most significantly enriched pathways in KOKI mice were *interferon-mediated signaling* and *regulation of inflammasome signaling*, whereas pathways related to *epithelial-to-mesenchymal transition, macrophage activation, mesenchymal cell migration, smooth muscle cell differentiation, cell-matrix adhesion and its regulation* were negatively enriched in FN ASO treated KOKI mice **(Figure 4F and Table S4)**. Consistent with the attenuation of the fibrotic program, *TGF-β1 receptor signaling*, *TGF-β production*, *inflammatory response to wounding* and *collagen biosynthetic processes* were also significantly downregulated **(Figure 4F and Table S4)**.

**Figure 4.**
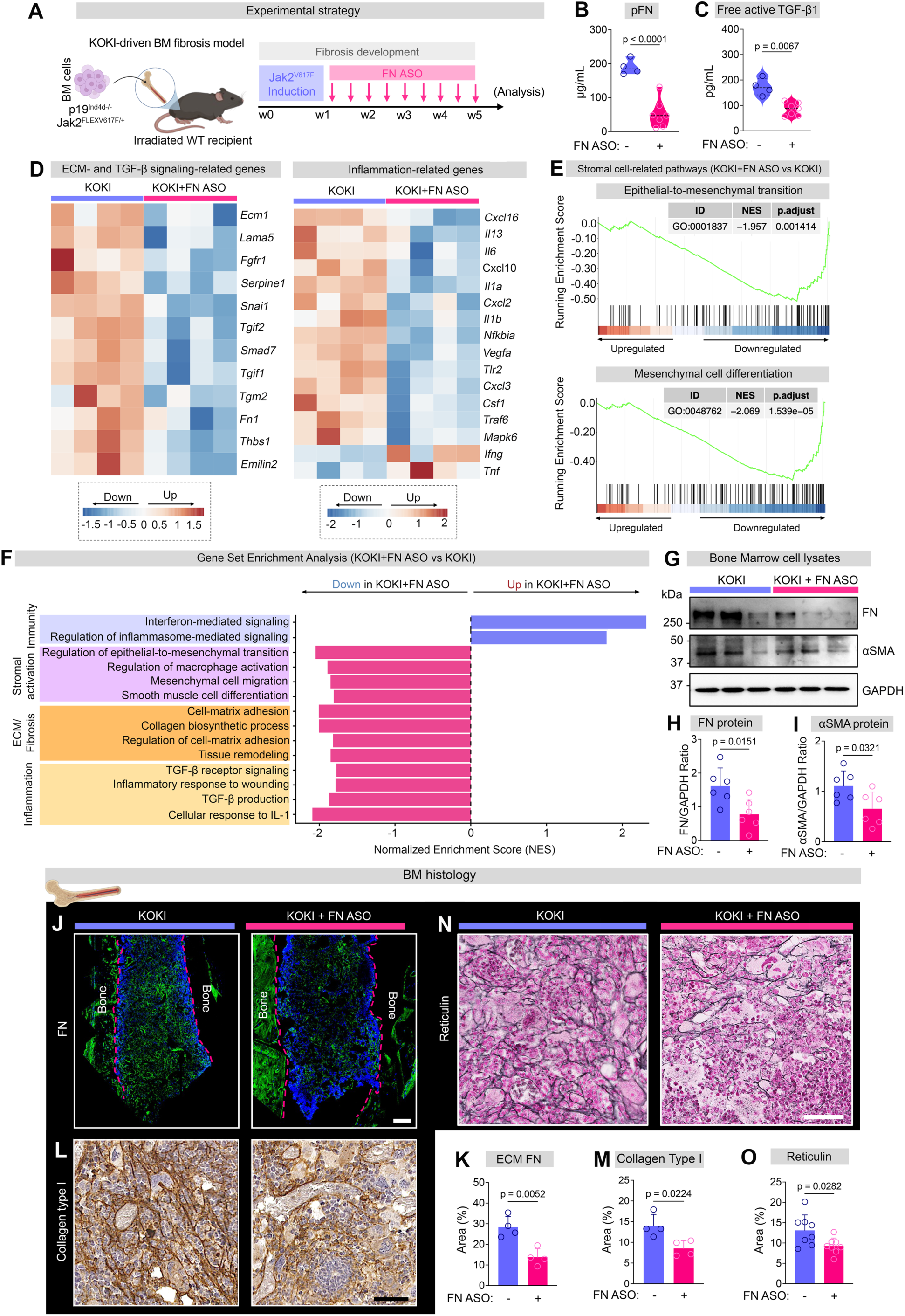
FN silencing remodels the fibrotic niche in *JAK2^V617F^*-driven myelofibrosis. **A)** Schematic representation of the experimental design for FN ASO or vehicle administration in KOKI mice. **B)** ELISA quantification of plasma FN levels in vehicle- (n = 5) and FN ASO-treated (n = 7) KOKI mice. Welch’s t test was used. **C)** ELISA quantification of active TGF-β1 level in BM cell-free fluids from vehicle- (n = 4) and FN ASO-treated (n = 7) KOKI mice. Welch’s t test was used. **D)** Heatmaps showing the average expression of ECM- and TGF-β signaling-related (left panel), and inflammation-related (right panel) genes in BM cells isolated from KOKI and KOKI plus FN ASO mice (n = 4). Color intensity represents scaled gene expression (z-score). **E-F)** Gene set enrichment analysis (GSEA) of selected differentially expressed genes in KOKI plus FN ASO versus KOKI mice. GSEA enrichment plots show the enrichment profiles of epithelial-to-mesenchymal transition and mesenchymal cell differentiation (E) pathways across the ranked gene list. Bar plots show significantly enriched pathways, with normalized enrichment scores (NES) indicating the direction and magnitude of enrichment (F). **G-I)** Representative western blot analysis of FN and α-SMA in BM cell lysates from vehicle- and FN ASO-treated KOKI mice (G), with corresponding relative quantification (H-I). At least four biological replicates were analyzed. GAPDH served as the loading control. Welch’s t test was used. **J-K)** Immunofluorescence staining of FN in BM sections from vehicle- (n = 3) and FN ASO-treated KOKI mice (n = 4) (J), with corresponding quantification (K). Nuclei (blue) were counterstained with Hoechst 33258. Images were acquired using a 25x objective; scale bar = 100 μm. Each data point represents one biological replicate and corresponds to the mean percentage of positive area measured across at least five independent fields. Welch’s t test was used. **L-O)** Representative immunochemical staining of type I collagen (L) and reticulin (N) in BM sections from vehicle- (n = 8) and FN ASO-treated (n = 8) KOKI mice, with corresponding quantification (M and O). Images were acquired using a 20x objective; scale bar = 100 μm. Each data point represents one biological replicate and corresponds to the mean percentage of positive area measured across at least five independent fields. Welch’s t test was used.

Collectively, these data indicate that FN silencing induces coordinated remodeling of the fibrotic BM microenvironment, characterized not only by attenuation of ECM/collagen and TGF-β1-associated programs, but also by a qualitative reshaping of inflammatory signaling and reduced stromal activation and mesenchymal remodeling. Reduced FN and α-SMA expression in the BM of FN ASO-treated mice was confirmed at the protein level **(Figure 4G-I)**. KOKI mice displayed extensive accumulation of elongated FN fibrillar networks in the BM, whereas FN ASO-treated mice exhibited markedly reduced microfibrillar FN organization **(Figure 4J-K)**. Consistently, histochemical BM analysis revealed decreased type I collagen **(Figure 4L-M)** and reticulin **(Figure 4N-O)** deposition after 5 weeks of FN ASO treatment, indicating reduced fibrotic ECM accumulation. Hematological analysis revealed only minor changes, with a significant increase in platelet counts **(Figure S4E)**. No changes in the BM frequencies of erythroid cells or megakaryocytes were detected **(Figure S4F)**; however, megakaryocytes exhibited a reduced cell size **(Figure S4G-H)**.

Similar to the findings in Romiplostim-treated mice, FN ASO treatment did not reduce the spleen-to-body weight ratio **(Figure S4I-J)** or affect extramedullary hematopoiesis **(Figure S4K)**. Nevertheless, FN ASO reduced FN and stromal α-SMA and collagen expression in the spleen **(Figure S4L-M)**, together with reduced FN matrix organization and reticulin deposition **(Figure S4N-O)**. Overall, these findings suggest that systemic FN reduction attenuates the fibrotic phenotype in a *Jak2*^V617F^-induced myelofibrosis mouse model.

## Discussion

Fibrotic remodeling of the BM niche is a key pathogenic feature of PMF and represents a major barrier to effective disease modification^21,22^. Although current therapies can effectively control splenomegaly and disease-related symptoms, directly targeting the ECM that sustains the fibrotic microenvironment remains challenging^23^. Lysyl oxidase inhibitors, which target collagen cross-linking, are currently under clinical evaluation; however, this approach primarily targets ECM stabilization rather than the upstream mechanisms that sustain pathological ECM production^24^.

Here, we identify FN as a functional component of this pathological niche and show that systemic FN silencing attenuates BM fibrosis in two complementary mouse models. Rather than acting solely as a structural component deposited during fibrosis, our findings support a role for FN in sustaining stromal activation, TGF-β1 activity, and the organization of the fibrotic matrix.

FN is particularly positioned to amplify fibrotic remodeling because it integrates matrix assembly with cell adhesion and growth factor signaling^25–28^. In MSCs, FN depletion suppressed gene programs associated with stromal cell activation and ECM production and remodeling, thereby impairing the TGF-β1-driven activated stromal phenotype. Consistent with these observations, systemic FN depletion *in vivo* reduced active TGF-β1 level, α-SMA and LTBP1 expression, reticulin, and collagen deposition in fibrotic BM. Collectively, these findings are consistent with a model in which FN matrix assembly contributes to a feed-forward circuit linking ECM organization to TGF-β1-dependent stromal activation.

We further show that circulating FN substantially contributes to BM ECM accumulation under fibrotic conditions, complementing the profibrotic activity of EDA-containing FN isoforms produced by resident stromal cells^16^. This distinction may be therapeutically relevant, as targeting FN expression systemically has the potential to affect both sources simultaneously. However, because the ASO used here does not discriminate between plasma- and stromal-derived FN, the relative contribution of each pool to disease progression remains unresolved and will require source-specific genetic or pharmacological approaches.

The effects of FN depletion were most pronounced on the stromal and ECM compartments, whereas correction of hematopoietic abnormalities was more variable across models. In Romiplostim-treated mice, FN silencing attenuated thrombocytosis and megakaryocyte expansion, consistent with previous evidence linking FN-dependent signaling to megakaryopoiesis^16,29^. Whether these effects arise through integrin-dependent regulation of progenitor fate, altered niche retention or disruption of aberrant stromal support remains to be determined.

By contrast, in the J*ak2*^V617F^-driven model, changes in hematopoietic output were comparatively modest despite a clear reduction in fibrotic matrix deposition and stromal activation. This divergence suggests that remodeling the extracellular niche may be insufficient to overcome hematopoietic phenotypes that are strongly driven by cell-intrinsic oncogenic signaling. It also supports the possibility that FN-directed therapy could be complementary, rather than alternative, to approaches targeting the malignant hematopoietic clone.

The transcriptional changes induced by *Fn1* silencing provide further insight into how FN contributes to the fibrotic BM niche. GSEA revealed a broad attenuation of stromal and mesenchymal programs in FN ASO-treated KOKI mice, including cell-matrix adhesion, mesenchymal cell migration, smooth muscle cell differentiation and TGF-β1 signaling, together with reduced collagen biosynthesis and inflammatory responses associated with tissue remodeling. These findings suggest that FN depletion may not simply reduce ECM accumulation, but may actively interfere with the stromal activation state that sustains fibrotic remodeling. Interestingly, interferon-associated pathways were relatively enriched following *Fn1* silencing, consistent with a qualitative reprogramming rather than a generalized suppression of inflammatory signaling. Whether these transcriptional changes reflect altered immune cell composition, changes in the activation state of resident myeloid populations, or indirect consequences of ECM remodeling remains to be established.

Together with the reduction in active TGF-β1, stromal α-SMA expression, and ECM deposition observed *in vivo*, these findings support a model in which FN may represent a functional link between matrix assembly, stromal activation, TGF-β1 activity, and inflammatory signaling.

Collectively, our findings identify FN as an active organizer of the fibrotic BM microenvironment and establish systemic FN silencing as a potential antifibrotic strategy in MF. By targeting a component of the pathological niche rather than the hematopoietic clone itself, FN ASO may provide a complementary therapeutic framework for MF and potentially extend to other fibrotic disorders in which pathological FN accumulation and ECM remodeling contribute to progressive organ fibrosis.

### Limitations of the study

This study has several limitations that warrant consideration. First, FN-targeting ASO demonstrated robust antifibrotic activity within a relatively short time frame, its effects on long-term disease progression and overall survival were not assessed. Second, FN is thought to play a prominent role in the early stages of fibrotic development, and accordingly, our approach appears to primarily prevent ECM deposition rather than promote the resolution of established BM fibrotic tissue. This highlights the importance of assessing its efficacy in models of pre-existing fibrosis. Third, our findings do not establish whether the reduction in BM fibrosis primarily results from altered megakaryocyte function, a direct effect of FN depletion on stromal cells, or the coordinated modulation of both compartments. Dissecting the relative contribution of these mechanisms will require cell-specific approaches to determine how FN regulates megakaryocyte-stromal cell crosstalk during myelofibrosis progression. Together, these limitations underscore the need for further studies to validate the translational potential of FN-targeting ASO therapy in clinical settings.

## Resource Availability

Human raw RNA-seq and mouse raw RNA-seq have been deposited in GEO under accession numbers *GSE350819* and *GSE350678*, respectively. Further information and requests for resources and reagents should be directed to, and will be fulfilled by, the lead contact, Alessandro Malara.

## Acknowledgements

We thank the animal facility and the OPBA of the University of Pavia for hosting the animals and supporting in animal protocol drawing up, and the Centro Grandi Strumenti of the University of Pavia for technical assistance with confocal microscopy, flow cytometry and Micro-CT imaging. This paper was supported by Associazione Italiana per la Ricerca sul Cancro (AIRC MFAG 2020 - project number 24541, A.M.), and Italian Ministry of University and Research (PRIN 2022 - project number 2022 AHXBBC_002; A.M.). The research activities of HR, CM and IP are supported by grants from Ligue Nationale Contre le Cancer (Equipe labellisée, 2022 and 2026 to HR).

## Author Contributions

F.R.C. designed, performed, and analyzed in vitro experiments with mesenchymal stromal cells and in vivo experiments in mice and wrote the manuscript. V.A. performed and analyzed experiments in mice. M.F.T. performed and analyzed experiments using hepatic cell lines. D.T. and U.G. performed histological analyses and interpreted the data. Carla Martinelli performed electron microscopy analyses and interpreted the results. S.D. performed bioinformatic analyses. Carolyne Marty, H.R., and I.P. provided transplanted KOKI mice and interpreted the experimental results. D.B. and M.P. designed the antisense oligonucleotide, contributed to project planning, designed and performed experiments, and analyzed and interpreted the results. A.B. contributed to project planning and analyzed and interpreted the results. A.M. secured funding, designed and performed experiments, analyzed and interpreted the results, and wrote the manuscript.

## Declaration of generative AI and AI-assisted technologies in the writing process

During the preparation of this work, the authors used OpenAI ChatGPT to improve language clarity and grammar. The authors reviewed and edited the content as necessary following the use of this tool and take full responsibility for the content of the publication.

## Material and Methods

### Animal studies

#### FN ASO treatment in naïve-wild type mice and in a Romiplostim-induced model of BM fibrosis

All animal experiments were reviewed and approved by the Italian Ministry of Health (approval no. 75/2022-PR and 498/2025-PR) and conducted in accordance with European regulations for animal experimentation. Wild-type C57BL/6J mice were obtained from Charles River Laboratories (Calco, Italy). Male and female mice aged 6–8 weeks were included in the experiments to minimize potential sex-related bias. All mice were maintained under a 12 h light/dark cycle, with ad libitum access to water and standard chow. Mice were randomly assigned to experimental groups. To induce BM fibrosis, mice received two subcutaneous injections of Romiplostim (1 mg/kg body weight; MedChem Tronica) at 7-day intervals (day 0 and 7) and sacrificed at day 14. In selected experiments, mice received five subcutaneous injections (day 0, 3, 7, 10 and 13) of Romiplostim at 100μg/kg before sacrifice at day 14. At day 14, 1 mg of FN or bovine serum albumin (Sigma-Aldrich) were labelled with NHS-Rhodamine (Thermo Fisher) and administered three times (0.33 mg/each) to vehicle- or Romiplostim-treated mice at 4h intervals before sacrifice. FN ASO was administrated five times (day 0, 3, 7, 10 and 13) over two weeks in naïve WT mice and nine times (day 0, 3, 7, 10, 14, 17, 21, 24, 26) over four weeks in Romiplostim-treated mice by subcutaneous injections at 50 mg/kg concentration. Control mice were injected with vehicle (Phosphate Buffer Saline, PBS).

### KOKI mice studies

KOKI mice were generated by crossing homozygous *p19*^Ink4d⁻/⁻^ KO mice with conditional *Jak2*^FLEXV617F/+;^ *SCL-Cre*^+/−^ KI mice, as previously described^30^. BM cells from non-induced KOKI mice (3 × 10^6^ cells; 15 × 10^6^ cells/mL in PBS) were transplanted into lethally irradiated (9.5 Gy; XRAD 320 kV irradiator) WT female recipient mice by retro-orbital injection. Four weeks after engraftment, *Jak2^V617F^* expression was induced with tamoxifen. Tamoxifen (Sigma-Aldrich) was dissolved in corn oil and administered by oral gavage at 200 mg/kg for five consecutive days. Mice were also maintained on tamoxifen-containing chow (400 mg/kg tamoxifen; Ssniff, Germany) during this week. FN ASO was administrated nine times (day 0, 3, 7, 10, 14, 17, 21, 24, 26) between weeks 2 and 5 post-*Jak2^V617F^* induction by subcutaneous injections at 50 mg/kg concentration. Control KOKI mice were injected with vehicle (PBS).

#### Antisense Oligonucleotide

The gapmer ASO has a DNA-like central region that directs cleavage of the complementary mRNA or pre-mRNA target by endogenous RNase H, and chemically modified wings that promote tighter RNA binding, enhanced stability, and improved cellular uptake. The FN1 mRNA target sequence was selected for high human-mouse conservation, enabling translational validation in preclinical models. It is positioned away from splice-regulatory elements (donor/acceptor sites and splicing enhancers/silencers) to ensure RNase H-mediated degradation without altering splicing. Secondary-structure prediction identified a single-stranded, accessible loop region to promote efficient Gapmer hybridization. Bioinformatic screening across human and mouse transcriptomes confirmed sequence uniqueness, minimizing off-target binding and enhancing safety. FN antisense oligonucleotide (FN ASO) (sequence 5’-GCUGAGCCTGCCTCTUGCUC-3’) and scramble ASO (sequence 5’-GCGACTATACGCGCAATATG-3’) were synthesized by Integrated DNA Technologies (Iowa, USA).

#### Blood collection and analyses

Blood was collected from the inferior vena cava using insulin syringes containing acid– citrate-dextrose (ACD) as anticoagulant (1 mL; BD Micro-Fine, 23G needle, 0.33 mm). Hematological parameters were analyzed using a Sysmex XN-1000Vet hematology analyzer (Sysmex, Japan) at the Istituto Zooprofilattico Sperimentale della Lombardia e dell’Emilia-Romagna (IZSLER, Brescia, Italy).

Platelet activation was assessed in diluted platelet-rich plasma (PRP). Blood was collected from the inferior vena cava after CO₂ euthanasia using acid-citrate-dextrose (ACD) as anticoagulant. Whole blood was centrifuged at 200 × g for 7 min at room temperature to obtain PRP. PRP was then diluted in Tyrode’s buffer supplemented with CaCl₂ to achieve a final calcium concentration of 1 mM and adjusted to a final platelet concentration of 2 × 10⁷ platelets/mL. Diluted PRP samples were stimulated with ADP (10 μM) or type I collagen (10 μg/mL) in the presence of fluorochrome-conjugated antibodies against activated integrin αIIbβ3 (JON/A, Emfret Analytics, Germany) or P-selectin/CD62P (Clone RMP-1, Biolegend). Samples were incubated for 20 min in the dark at room temperature and then fixed with 0.2% formalin saline. Platelet activation was analyzed by flow cytometry.

Platelet-poor plasma (PPP) was prepared by centrifuging PRP at 600g for an additional 13 min. The supernatant was collected, clarified by centrifugation at 10,000 rpm for 10 min, and stored at -80 °C until use. BM cell-free fluids were obtained by flushing femurs and tibias with 1 ml PBS, followed by centrifugation at 13,000 rpm for 20 min. The resulting supernatant was collected and stored at -80 °C until analysis.

#### Tissue Specimens

Bones, livers and spleens from mice were recovered from mice and fixed in buffered formalin or in 4% paraformaldehyde solutions. Bones were decalcified in EDTA-based MicroDec (Diapath, Italy) at 4°C. Gomori staining was performed on paraffin embedded BM sections using Reticulum II staining kit (Ventana; Roche) according to the manufacturer’s instructions.

#### Cell cultures

hMSCs were purchased from Lonza. All experiments were performed using early passage hMSCs, maximum passage 6. hMSCs were grown in 12-well plate to 70-80% confluence in high-glucose DMEM (Euroclone, Italy), supplemented with 1% of penicillin/streptomycin (Euroclone), 1% of L-glutamine (Euroclone), 10% of MesenCult (Stem Cell Technologies, Vancouver, CA), and 1% of non-essential aminoacids (NEAAs; Euroclone, Italy). C57BL/6 mouse embryonic fibroblasts (mEFs) were from ATCC and cultured in DMEM (Euroclone, Italy) and 10% Fetal Bovine Serum (FBS) (Gibco BRL).

#### In vitro ASO transfection

hMSCs and mEFs were transfected with 150 nM of FN ASO or scramble ASO using Lipofectamine RNAiMax (Invitrogen, Italy). Briefly, the proper amount of Lipofectamine RNAiMax and FN ASO or scramble ASO were added to serum-reduced Opti-MEM (Gibco BRL) and allowed to complex for 5 min at room temperature. Then, the complex was supplied to the cells in the presence or absence of 10 ng/mL of human recombinant TGF-

β1 (Peprotech, UK) and incubated at 37°C and 5% CO_2_ for 24 h. For dose-response experiments, FN ASO was also used at 12, 250 and 500 nM concentrations. Cultured Huh-7 and Hepa 1-6 cell lines, were transfected using 3 μL LipofectAMINE2000®/ml OptiMEM with escalating dose of 0.015-1 µM ASOs. Following approximately 24 h of treatment, the medium was replaced with fresh medium, and the cells were cultured overnight. Total RNA was then isolated, and *FN1* mRNA expression was quantified by qRT-PCR.

#### TLR9 activity assay

HEK-Blue™ mouse TLR9 reporter cell lines (InvivoGen, Milan, Italy) are stable HEK293-derived cells co-expressing the TLR9 gene and a secreted embryonic alkaline phosphatase (SEAP) reporter gene. The SEAP reporter is under the control of an IFN-β promoter fused to NF-κB and AP-1 binding sites, enabling SEAP production upon TLR9 activation. HEK-Blue™ mTLR9 cells were cultured according to the manufacturer’s instructions in high-glucose DMEM supplemented with 10% (v/v) FBS, 100 U/mL penicillin, 100 μg/mL streptomycin, 2 mM L-glutamine and 100 μg/mL Normocin, and maintained at 37 °C in a humidified 5% CO₂ incubator. All experiments were performed using early-passage cells (≤6 passages). Cells were seeded at a density of 1 × 10^5^ cells per well in 96-well plates containing 180 μL HEK-Blue™ Detection Medium (InvivoGen). Cells were stimulated in triplicate with 20 μL of the TLR9 agonist ODN1668 (1 μM; InvivoGen), the inhibitor ODN2088 (10 μM; InvivoGen), FN ASO (150 nM), or PBS as a negative control, and incubated for 16 h. SEAP activity was quantified by measuring the optical density at 620 nm using a microplate spectrophotometer (HiPo MPP-96, Biosan, Latvia).

#### In vitro Wound healing assay

For wound healing assays, hMSCs were cultured in complete medium in 6-well plates until 100% confluence and transfected as described above. A linear scratch (“wound”) was generated across the cell monolayer using a sterile pipette tip. Cells were then incubated with fresh complete medium alone or supplemented with recombinant human TGF-β1 (10 ng/mL). Images were acquired immediately after scratching (T_0_) and after 24 h (T_1_) using an IX53 Olympus microscope equipped with a UPlanFL 4×/0.13 objective. The area covered by migrating cells was quantified using ImageJ software (NIH, USA).

#### Collagen gel contraction assay

For collagen gel contraction assays, hMSCs were transfected overnight in 6-well plates as described above. Cells were then detached using trypsin-EDTA (Euroclone) and embedded in three-dimensional collagen gels (3 mg/mL) prepared from rat tail collagen type I (Ibidi GmbH, Gräfelfing, Germany) in the presence of scramble (150 nM), FN ASO (150 nM) and/or recombinant human TGF-β1 (10 ng/mL, Peprotech, UK). Gels were allowed to polymerize for 1 h at 37 °C in a humidified incubator with 5% CO₂. After detachment from the well edges, collagen gels were overlaid with complete culture medium containing scramble, FN ASO and/or TGF-β1. Medium was replaced every 2 days for 10 days. Gel contraction was quantified using ImageJ software.

#### Immunofluorescence and immunohistochemistry

hMSCs were seeded on 13-mm glass coverslips in 24-well plates, transfected as described above, and fixed after 24 h with 4% paraformaldehyde for 20 min. Cells were washed with PBS and permeabilized with 0.1% Triton X-100. FN was detected by overnight incubation at 4 °C with a rabbit anti-FN primary antibody (1:250; Millipore), followed by incubation with a goat anti-rabbit Alexa Fluor 488 secondary antibody (1:500; Invitrogen) for 1 h at room temperature. Nuclei were stained with Hoechst 33258 (100 ng/mL in PBS). Coverslips were mounted on glass slides using ProLong Gold antifade reagent (Invitrogen).

For tissue staining, OCT-embedded BM, liver and spleen sections were incubated overnight at 4 °C with the rabbit anti-FN (1:100), or anti-phosphorotioate (1:100; clone PS03, Rockland) antibodies, followed by goat anti-rabbit Alexa Fluor 488 secondary antibody (1:500) for 4 h at room temperature. Images were acquired using Nikon Eclipse Ti2 fluorescence microscope (Nikon) equipped with a Plan Apo 10×/0.45 objective or a DMi8 SP8 confocal laser scanning microscope using a 25x water-immersion objective (HC FLUOTAR L 25x/0.95 W DLS, Leica Microsystems).

For immunohistochemistry, buffered formalin-fixed and decalcified femurs were dehydrated through 70% ethanol (60 min), two changes of 95% ethanol (90 min each), and three changes of 99% ethanol (60 min each), followed by clearing in three changes of xylene (90 min each). Samples were then immersed in three changes of paraffin (1 h each), embedded in paraffin blocks, and stored at room temperature until sectioning. Hematoxylin and eosin (H&E) staining (Diapath) was performed on serial sections to assess tissue morphology. For immunohistochemical analysis, paraffin was removed with xylene, and sections were rehydrated through graded alcohols. Antigen retrieval was performed using preheated Target Retrieval Solution, pH 6.0, for 30 min. Sections were then blocked with fetal bovine serum in PBS for 60 min and incubated overnight with a rabbit monoclonal anti-type I collagen antibody (1:200; clone E8F4L; Cell Signaling Technology). Antibody binding was detected using a donkey anti-mouse IgG (whole molecule)–peroxidase antibody produced in rabbit (1:200; Sigma, cat. no. A9044) for 2 h, followed by visualization with a diaminobenzidine (DAB) chromogen reaction (Peroxidase Substrate Kit, SK-4100; Vector Laboratories). Sections were counterstained with Mayer’s hematoxylin. Histological image quantification of FN, reticulin, and type I collagen deposition were quantified using ImageJ2 (NIH, v.2.16.0). Analyses and scoring were performed blinded to treatment group. BM regions of interest (ROIs) were defined by excluding cortical and trabecular bone and other non-marrow structures. Images were converted to binary masks using a predefined threshold for each staining, which was applied consistently across samples within the same experiment. The percentage of positively stained area was calculated relative to the total BM ROI. At least 5 fields from 3 sections were analyzed for each animal and averaged to obtain a single value per mouse. Individual mice were considered independent biological replicates for statistical analyses. For Megakaryocytes density and size quantification, femurs were scanned and analyzed with Nano Zoomer Digital Pathology View2 (Hamamatsu). For megakaryocyte density quantification, BM sections were divided into five regions of approximately equivalent size for each mouse. The area of each region was measured, and the number of megakaryocytes within each region was manually counted. Megakaryocyte density was calculated for each region as the number of megakaryocytes divided by the corresponding analyzed area. The five regional measurements were then averaged to obtain a single megakaryocyte density value for each mouse. Data from individual mice were used as biological replicates for statistical analysis. For megakaryocyte size quantification, megakaryocytes were randomly selected throughout the entire BM section, without restricting the analysis to specific anatomical or histological regions, including areas adjacent to the bone or regions displaying different degrees of fibrosis. This sampling strategy was used to minimize selection bias and to provide a representative assessment of megakaryocytes size across the BM section. A minimum of 25 megakaryocytes were analyzed per mouse. Measurements were averaged within each mouse to generate a single biological replicate for statistical analysis.

#### Transmission Electron Microscopy (TEM)

In order to investigate structural and ultrastructural features, samples were fixed in 2.5% glutaraldehyde in 0.13 M phosphate buffer pH 7.2-7.4 for 2 hours, post-fixed in 1% osmium tetroxide, dehydrated through graded ethanol and propylen oxide and embedded in epoxy resin. Several semithin sections were prepared from each sample and stained with 0.5% toluidine blue in 1% sodium borate. Ultrathin sections of 50 to 60 nm were counterstained with Pt-blue and lead citrate, and observed using a Jeol JEM 1010 transmission electron microscope (Jeol, Tokyo, Japan).

### Micro-computed tomography (µCT) analysis

Right tibias from vehicle and FN ASO-treated mice were dissected, cleared of surrounding muscle and connective tissue, fixed in 4% paraformaldehyde (PFA) overnight at 4 °C, and stored in phosphate-buffered saline (PBS) prior to high-resolution scanning to assess cortical and trabecular bone microarchitecture. For µCT acquisitions, samples were wrapped in PBS-moistened gauze, sealed in Parafilm to prevent dehydration, and secured in a sample holder. Scans were acquired using a SkyScan 1276 µCT scanner (Bruker, Kontich, Belgium) with the following settings: source voltage 55 kV, source current 72 µA, 0.25 mm aluminum filter, 1×1 camera binning, exposure time 1750 ms, 3-frame averaging, nominal pixel size 4 µm, rotation step of 0.2° over a 180° rotation. Image reconstruction was performed using SkyScan NRecon software (v. 2.2.0.6), and 3D quantitative morphometric analysis was conducted using CT Analyser (CTAn v. 1.20.8.0, Bruker). The proximal growth plate was set as the reference slice. Trabecular and cortical bone analyses were performed in 1.0 mm and 0.8 mm high regions, located 0.5 mm (proximal metaphysis) and 5.0 mm (mid-diaphysis) distal to the growth plate, respectively. According to standard guidelines (Bouxsein et al., 2010), 3D analysis was used to determine trabecular bone volume fraction (BV/TV, %), thickness (Tb.Th, mm), separation (Tb.Sp, mm), and number (Tb.N, mm-1), as well as cortical BV/TV (%) and thickness (Ct.Th, mm). Bone mineral density was calibrated using two hydroxyapatite (HA) phantom rods (0.25 and 0.75 g HA/cm3, 2 mm diameter) scanned under identical conditions. Tissue mineral density (TMD, g HA/cm3) was reported for cortical bone, whereas bone mineral density (BMD, g HA/cm3) was reported for trabecular bone. 3D visual representations of representative samples were generated using CTvox software (v. 3.3.1, Bruker).

#### Bulk RNA sequencing

Nucleic acid extraction and purification were performed by RNA Extraction Kit (Qiagen). RNA samples were quantified, and their integrity (RIN, RNA integrity number) was assessed using High Sensitivity RNA ScreenTape Assay on 4200 TapeStation System (Agilent Technologies). 500 pg RNA was used from each sample for cDNA generation using the Takara Smart-Seq v4 Ultra Low Input RNA kit (Takara Bio USA, Ann Arbor, MI) following the manufacturer’s instructions, with 15 cycles of cDNA amplification. This technology relies on the template-switching activity of the reverse transcriptase to enrich for full-length cDNAs and to add defined PCR adapters directly to both ends of the first-strand cDNA. This ensures that the final cDNA libraries incorporate the 5′ end of the mRNA and maintain a true representation of the original mRNA transcripts. Smart-Seq cDNA was assessed for quality on High Sensitivity D5000 Screen Tape Assay on 4200 TapeStation System. Starting from 500 pg of cDNA, libraries were prepared through DNA library prep Illumina kit (Illumina, San Diego, CA, USA), following manufacturer’s instructions. This protocol is based by tagmentation drive on transposase cutting the genomic DNA and attaching the adapters on each side. Final libraries were quantified (High Sensitivity D5000 Screen Tape Assay) and pooled at 0,8nM concentration, then sequenced 1x100bp on the Illumina NovaSeq 6000 platform. For the hMSC experiment, single-end reads (100 bp) were trimmed with Trimmomatic (v0.39) to remove adapters and exclude low-quality reads from the analysis and aligned to the GRCh38 reference genome using STAR aligner (v2.5.3a). FeatureCounts (v1.6.4) was used to compute reads mapping on exons as annotated in GENCODE Human basic gene annotation (version 31) and obtain transcript quantification summarized at the gene level, excluding multimapping reads. Only genes showing an average expression greater than 1 CPM (Counts per million) were retained. Differential gene expression analysis was performed with DESeq2 Bioconductor library (v1.38.3). Heatmaps were made with Pheatmap R package (v1.0.12). GO analysis was conducted with Enrichr R library (v3.1) on DEGs (|log2FC|>0.5, adj.p-value<0.05). For the KOKI mouse samples, paired-end reads (100 bp) were trimmed with Trimmomatic (v0.38) to remove adapters and exclude low-quality reads from the analysis and aligned to the Mus musculus reference genome (mm10) using HISAT2 aligner (v2.2.1). StringTie (v2.1.3b) was used to compute reads mapping on exons as annotated in NCBI annotation (version 108) and obtain transcript quantification summarized at the gene level. Only genes showing an average expression greater than 1 CPM (Counts per million) were retained. Differential gene expression analysis was performed with DESeq2 Bioconductor library (v1.52.0). Heatmaps were made with Pheatmap R package (v1.0.13). GO analysis was conducted with Enrichr R library (v3.4) on DEGs (|log2FC|>0.5, adj.p-value<0.05). Gene set enrichment analysis was performed using all expressed genes (cpm > 1) as input, with the clusterProfiler package (v4.20.0).

#### qPCR and splicing assay

RNA from cells and tissue was extracted with RNeasy Plus Micro Kit (Qiagen), according to the manufacturer’s instructions. 15 μL of total RNA was retrotranscribed into complementary DNA (cDNA) using the iScript cDNA Synthesis Kit (Bio-Rad Laboratories, Italy). The conditions for the reverse transcription were as follows: 25°C for 5 min, 46°C for 20 min, 95°C for 1 min. FN splicing assay was performed as previously described^31,32^. Briefly, 300 ng and 500 ng of cDNA derived from mEFs and mouse tissues, respectively, were amplified by PCR in reactions containing 10x MgCl_2_-supplemented Buffer I, deoxynucleoside triphosphates (dNTPs; 0.5 mM; Invitrogen), primers (0.2 μM; Sigma), and AmpliTaq™ DNA Polymerase (0.625 U; Applied Biosystems). Amplification was performed under the following cycling conditions: 95°C for 2 min, followed by 35 cycles of 95°C for 15 sec, 55.4°C (*EDA*) or 46°C (*IIICS*) for 30 sec, 72°C for 1 min, with a final extension of 72°C for 5 min. Sample were resolved on a 3% agarose gel in TBE solution and visualized using the ChemiDoc™ Touch Image System (Bio-Rad).

For quantitative Real-Time PCR, RT samples were diluted up to 60 µL with ddH_2_O and 3 µL of the resulting cDNA was amplified in triplicate in 20 µL reaction mixture with 125 nM of each specific primer and SsoFast™ EvaGreen® Supermix (Biorad Laboratories, Milan, Italy). The amplification reaction was performed in a CFX Real-time system (BioRad Laboratories) as follows: 95°C for 30 sec, followed by 35 cycles at 95°C for 15 sec, 58°C for 30 sec, 70°C for 15 sec and 95°C for 5 sec. The following pre-designated KiCqStart™ primers for *Fn1*, *Col3a1*, *Acta2*, *Il6* and *Tgfb1* genes were purchased from Sigma-Aldrich. GAPDH and β2m gene expression were used for the comparative concentration analysis.

#### Western Blot

Cells were lysed in Hepes-glycerol lysis buffer (50 mM Hepes, 150 mM NaCl, 10% glycerol, 1% Triton X-100, 1.5 mM MgCl_2_, and 1 mM EGTA) containing 1 µg/mL leupeptin and 1 µg/mL aprotinin for 30 min at 4°C. Samples were clarified by centrifugation at 13,000 rpm at 4°C for 15 min. 3X Laemmli sample buffer was then added to protein extracts. Samples were always heated at 95°C for 5 min, separated by electrophoresis on 10%, 12% or 15% sodium dodecyl sulfate-polyacrylamide gel, and then transferred to polyvinylidine fluoride membranes. Membranes were probed with primary antibodies, washed three times with PBS and Tween20 0.1% (Biorad Laboratories), and incubated with an appropriate peroxidase-conjugate secondary antibody (1:3000; Biorad Laboratories). Membranes were visualized using the ChemiDoc™ Touch Image System (Bio-Rad). The following antibodies were used: rabbit polyclonal anti-GAPDH (1:4000; ABS16, Millipore), mouse monoclonal anti-α-SMA (1:2000; clone 1A4, Sigma-Aldrich), rabbit polyclonal anti-Fibronectin (1:2000; AB1954, Millipore), rabbit polyclonal anti-LTBP1 (1:500; #26855-1-AP, Proteintech), rabbit polyclonal anti-TGF-β1 (1:1000; SAB4502954, Sigma-Aldrich), rabbit monoclonal anti-transferrin (1:1000; GTX101035, Genetex), rabbit monoclonal anti-transferrin receptor (1:1000; clone EPR20584, Abcam) and mouse monoclonal anti-Vimentin (1:1000; clone V9, Sigma-Aldrich). Western blot band intensities were quantified by densitometric analysis after background subtraction. The intensity of each target protein band was normalized to the corresponding GAPDH band in the same sample. The resulting target protein/GAPDH ratios were used for quantitative analysis and statistical comparisons between vehicle- and FN ASO-treated groups. The intensities of plasma FN and transferrin bands, as well as, TGF-β1 bands in BM cell-free fluids, were normalized to the corresponding albumin band detected by Coomassie blue staining.

#### Deoxycholate (DOC) fractionation

Cultured hMSCs were solubilized using 200 μL of deoxycholic acid (DOC) lysis buffer (DOC 2%, 50 mM EDTA (ethylenediaminetetraacetic acid), 2mM phenylmethanesulfonyl fluoride, 2mM Iodoacetamide, 2mM N-ethylmaleimide and protease inhibitors). After centrifugation, the DOC-insoluble pellet was solubilized in 25 μL of 2% SDS, 20mM Tris-HCl, pH 8.8, 2mM phenylmethanesulfonyl fluoride, 2mM Iodoacetamide, 2mM N-ethylmaleimide, and 2 mM EDTA. Samples were immunoblotted and incubated with rabbit polyclonal anti-Fibronectin (1:2000; AB1954, Millipore). Rabbit polyclonal anti-GAPDH (1:4000; ABS16, Millipore) and mouse monoclonal anti-Vimentin (1:1000; clone V9, Sigma-Aldrich) were used to ensure the equal amount of proteins in the DOC-soluble and DOC-insoluble fractions, respectively.

#### ELISA assays

Soluble plasma FN was quantified using the Mouse Fibronectin ELISA Kit (#108849; Abcam), according to the manufacturer’s instructions. The levels of free active TGF-β1 were quantified in BM cell-free fluids using the LEGEND MAX Free Active TGF-β1 ELISA Kit (BioLegend, USA), according to the manufacturer’s instructions. Absorbance was measured at 450 nm using a microplate spectrophotometer (HiPo MPP-96, Biosan, Latvia).

### Flow Cytometry

Cell populations from BM and spleen were immunophenotypically characterized by flow cytometry using a BD FACSLyric™ Flow Cytometer (BD Biosciences). BM and spleen cell suspensions were stained with fluorochrome-conjugated antibodies according to the manufacturer’s recommendations.

Erythroid maturation stages were analyzed in whole spleen and BM cell suspensions by flow cytometry based on CD71 and Ter119 expression. Cells were classified into four populations (R1-R4) corresponding to progressively more mature erythroid precursors, from early erythroid progenitors (R1) to late-stage erythroblasts and erythroid cells (R4). R1 proerythroblasts were identified as TER119^low^/CD71^high^, R2 basophilic erythroblasts as TER119^high^/CD71^high^, R3 late basophilic and polychromatophilic erythroblast as TER119^high^/CD71^med^, and R4 orthochromatophilic erythroblasts as TER119^high^/CD71^low^, respectively. For analysis of megakaryocyte, monocyte/granulocyte, and lymphoid populations, red blood cells were lysed using a buffer containing 150 mM ammonium chloride, 10 mM potassium bicarbonate, and 0.1 mM EDTA. Cells were then stained with the following antibodies: anti-mouse CD41-PE (clone MWReg30) and anti-mouse CD45-APC (clone 30-F11) for megakaryocytes; anti-mouse/human CD11b-PE (Mac-1, clone M1/70) and anti-mouse Ly6G-Ly6C-FITC (Gr-1, clone RB6-8C5) for monocytes/granulocytes; anti-mouse CD3ε-FITC (clone 145-2C11) for T cells; and anti-mouse/human CD45R-PE (B220, clone RA3-6B2) for B cells. All antibodies were purchased from BioLegend (UK).

Cell viability was assessed in hMSCs transfected with scramble control or FN ASO using the Annexin V-FITC apoptosis detection kit (#IM3546, Beckman Coulter), according to the manufacturer’s instructions. Briefly, cells were gently detached by scraping, washed three times with ice-cold PBS, and centrifuged at 500 g for 5 min at 4°C. Cells were resuspended in 100 µL of 1x binding buffer. Annexin V-FITC and propidium iodide (PI) were then added, and cells were incubated for 15 min on ice in the dark. Finally, 400 µL of ice-cold 1x binding buffer was added, and samples were immediately analyzed by flow cytometry.

#### Statistics

Results are displayed as mean ± standard deviation (SD) from at least three independent biological replicates per group, or as indicated. Biological replicates were defined as independent cell-culture experiments or individual animals. Technical replicates, multiple microscopic fields, and individual cells obtained from the same biological sample were not considered independent biological replicates. For histological and immunofluorescence analyses, measurements from multiple fields were averaged for each animal before group-level statistical analysis, unless otherwise specified. Comparisons between two independent groups were performed using two-tailed unpaired Welch’s t-test. Comparisons among three or more groups were performed using one-way ANOVA followed by Tukey’s multiple-comparisons test as specified in the corresponding figure legends. Experiments involving two independent factors were analyzed using two-way ANOVA, including assessment of the interaction between factors, followed by Šídák’s multiple-comparisons test where appropriate. Statistical significance was defined as P < 0.05. Data were analyzed and plotted using GraphPad Prism software v.11.1.0.

