## Supplemental Figures and Tables for "Fibronectin silencing remodels the fibrotic bone marrow niche"

#### Supplementary Figure 1

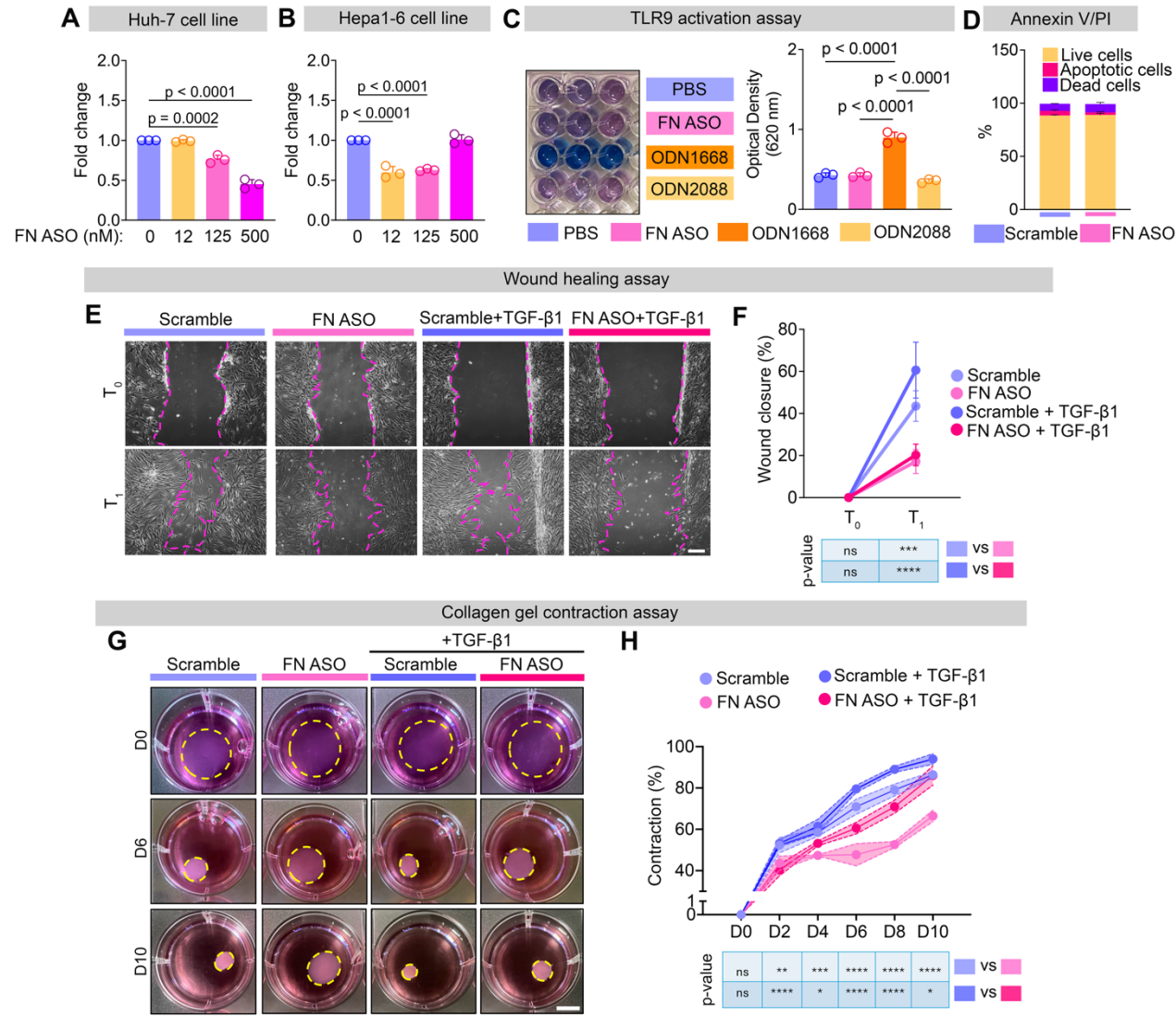

**Figure S1. Characterization of in vitro FN ASO specificity and safety.**

**A-B)** RT-qPCR analysis of FN1 expression in human Huh-7 (**A**) and murine Hepa1-6 (**B**) cells following treatment with scramble (0 nM) or FN ASO (12, 125 and 500 nM) for 24 hours (n=3). Ordinary one-way was used.

**C)** Assessment of TLR9 activation by FN ASO in HEK-Blue™ mouse TLR9 reporter cells. ODN1668 was used as a positive control, whereas PBS and ODN2088 were used as negative controls (n=3). Ordinary one-way was used.

**D)** Cell viability assay using Annexin V and propidium iodide (PI) staining in hMSCs treated with scramble or FN ASO (n=3). Welch's t test was used.

**E-F)** Wound healing assay assessing cell migration in hMSCs treated with scramble, FN ASO, TGF-β1 plus scramble or FN ASO (**E**) and quantification of wound closure (**F**). Objective = 4x. Scale bar = 50 μm. \*\*\*: $p < 0.001$ and \*\*\*\*: $p < 0.0001$. Data are representative of three independent experiments. Two-way Repeated Measures (RM) ANOVA was used.

**G-H)** Collagen gel contraction assay assessing cell contractility in hMSCs treated with scramble, FN ASO, TGF-β1 plus scramble or FN ASO (**G**) and quantification of gel contraction (**H**). Scale bar = 1 cm. \*: $p < 0.05$; \*\*: $p < 0.01$; \*\*\*: $p < 0.001$ and \*\*\*\*: $p < 0.0001$. Data are representative of three independent experiments. Two-way Repeated Measures (RM) ANOVA was used.

#### Supplementary Figure 2

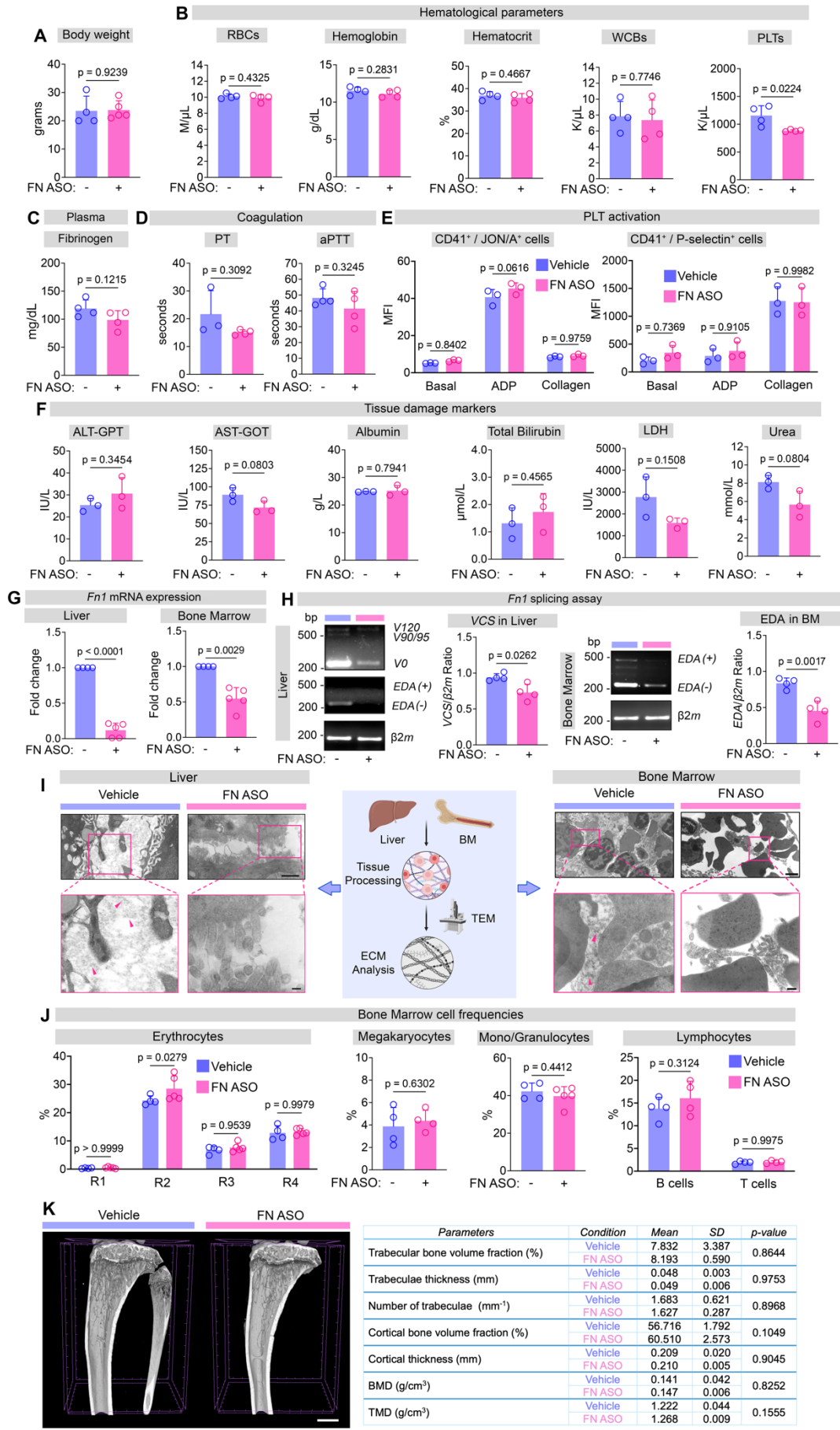

**Figure S2. FN ASO was well tolerated during short-term treatment *in vivo*.**

**A)** Body weight of vehicle- (n=4) and FN ASO-treated (n=5) mice at the end of the treatment window. Welch's t test was used.

**B)** Hematological parameters of vehicle- and FN ASO-treated mice (n=4), including red blood cell (RBC) count, hemoglobin, hematocrit, white blood cell (WBC) count and platelet (PLT) count. Welch's t test was used.

**C)** Quantification of fibrinogen levels in plasma samples from vehicle- and FN ASO-treated mice (n = 4). Welch's t test was used.

**D)** Prothrombin time (PT) and partial thromboplastin time (aPTT;) measured in plasma from vehicle- and FN ASO-treated mice (n = 3). Welch's t test was used.

**E)** Median fluorescent intensity (MFI) of CD41+/JON/A+ (J) CD41+/P-selectin+ (K) PLTs following ADP- and collagen-induced PLT activation in vehicle- and FN ASO-treated mice (n = 3). Ordinary two-way ANOVA was used.

**F)** Plasma levels of tissue damage markers in vehicle- and FN ASO-treated mice (n=3), including alanine aminotransferase (ALT-GPT), aspartate aminotransferase (AST-GOT), albumin, total bilirubin, lactate dehydrogenase (LDH) and urea. Welch's t test was used.

**G)** RT-qPCR analysis of *Fn1* expression in liver and BM cells from vehicle- and FN ASO-treated mice. At least 4 biological replicates were analyzed. Welch's t test was used.

**H)** PCR-splicing assay of EDA- and IIICS-containing FN isoforms in liver and BM from vehicle- and FN ASO-treated mice and their quantification. β2-microglobulin (β2m) was used as the loading control. mRNA levels were normalized to β2m. Welch's t test was used.

**I)** Transmission electron microscopy images of liver and BM samples from vehicle- and FN ASO-treated mice. Pink arrows indicate ECM structures. Scale bars = 500 nm (upper panels) and 150 nm (lower panels).

**J)** Cell frequencies of BM populations in vehicle- and FN ASO-treated mice, including erythrocytes, megakaryocytes, monocytes/granulocytes and lymphocytes (n = 4). R1= Proerythroblasts, R2= Basophilic erythroblasts, R3= Polychromatophilic erythroblasts, R4= Orthochromatophilic erythroblasts. Welch's t test was used for megakaryocytes and mono/granulocytes quantification; Ordinary two-way ANOVA was applied for erythrocytes and lymphocytes, respectively.

**K)** Representative μCT analysis of tibias from vehicle- and FN ASO-treated mice with quantification of trabecular bone volume fraction, trabeculae thickness, number of trabeculae, cortical bone volume fraction, cortical thickness, bone mineral density (BMD)

and tissue mineral density (TMD). Data are representative of three independent experiments. Welch's t test was used.

### Supplementary Figure 3

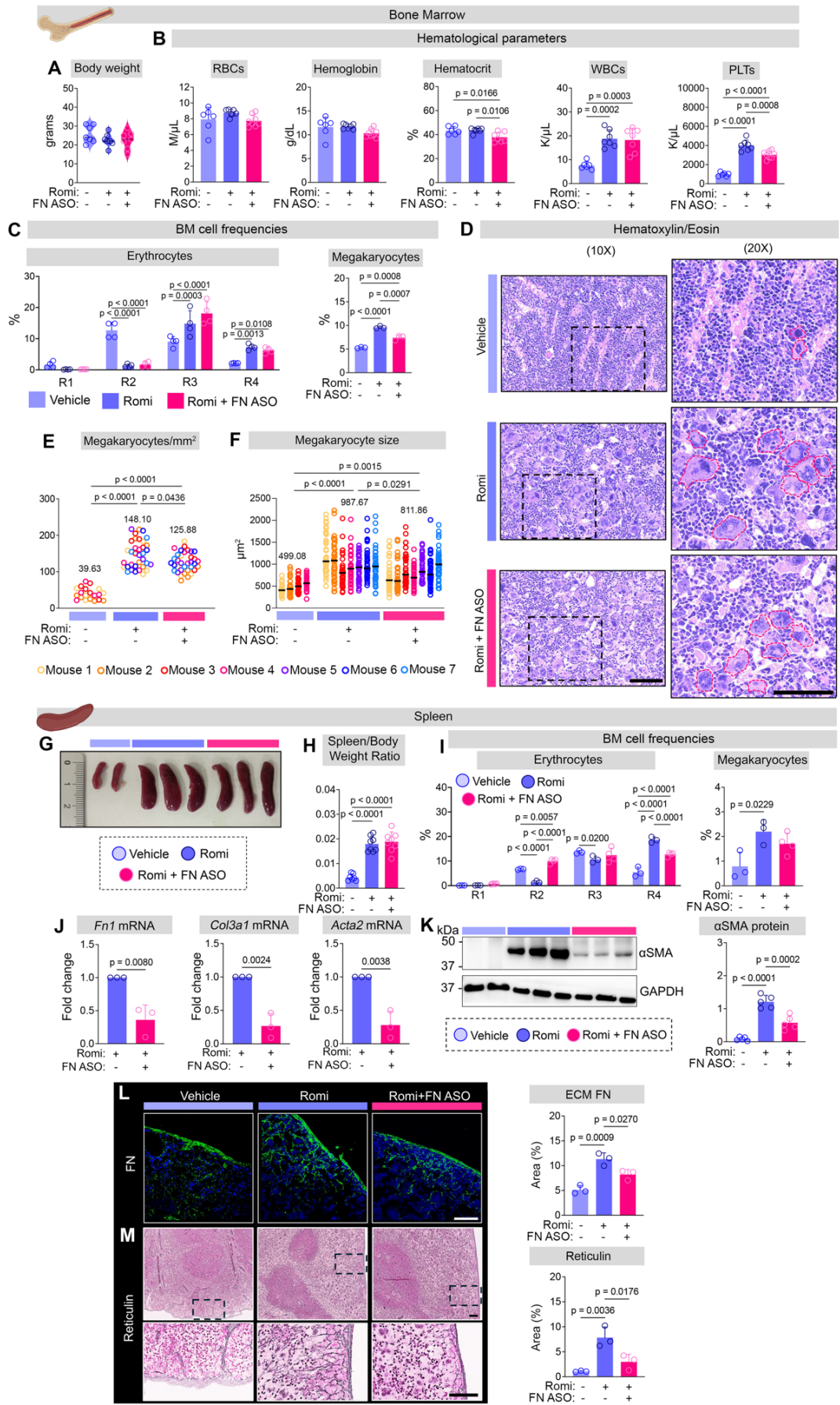

**Figure S3. Hematopoietic and splenic changes in fibrotic mice treated with FN ASO.**

**A)** Body weight of vehicle-, Romiplostim+vehicle- and Romiplostim+FN ASO-treated mice (n=7). Ordinary one-way ANOVA was used.

**B)** Hematological parameters of vehicle-, Romiplostim+vehicle- and Romiplostim+FN ASO-treated mice (n=7), including RBC count, hemoglobin, hematocrit, WBC and PLT counts. Ordinary one-way ANOVA was used.

**C)** Cell frequencies of BM erythrocytes and megakaryocytes in BM of vehicle-, Romiplostim+vehicle- and Romiplostim+FN ASO-treated mice. Data are representative of three biological replicates. R1= Proerythroblasts, R2= Basophilic erythroblasts, R3= Polychromatophilic erythroblasts, R4= Orthochromatophilic erythroblasts. Ordinary two-way ANOVA and ordinary one-way ANOVA were used, respectively.

**D-F)** Representative hematoxylin and eosin staining of BM sections from vehicle-, Romiplostim+vehicle- and Romiplostim+FN ASO-treated mice (**D**). Quantification of megakaryocyte density and size are provided in **E** and **F**, respectively. Scale bars=250 μm. Each data point represents the mean value calculated from multiple cells/fields per mouse, with each mouse considered one biological replicate. Ordinary one-way ANOVA was used.

**G-H)** Representative spleens from vehicle-, Romiplostim+vehicle- and Romiplostim+FN ASO-treated mice (**G**) with spleen weight normalized to body weight (**H**).

**I)** Cell frequencies of erythrocytes and megakaryocytes in spleens from vehicle-, Romiplostim+vehicle- and Romiplostim+FN ASO-treated mice (n = 3). Ordinary two-way ANOVA and ordinary one-way ANOVA were used, respectively.

**J)** RT-qPCR analysis of *Fn1*, *Col3a1* and *Acta2* in spleens from Romiplostim+vehicle- and Romiplostim+FN ASO-treated mice (n = 3). Welch's test was used.

**K)** Representative western blot analysis and relative quantification of α-SMA expression in spleen cells from vehicle-, Romiplostim+vehicle- and Romiplostim+FN ASO-treated mice. GAPDH was used as a loading control. Ordinary one-way ANOVA was used.

**L)** Immunofluorescence analysis of FN in spleen sections from vehicle-, Romiplostim+vehicle- and Romiplostim+FN ASO-treated mice. Objective: 25x, scale bar = 150 μm. Ordinary one-way ANOVA was used.

**M)** Representative reticulin staining in spleen sections from vehicle-, Romiplostim- and Romiplostim+FN ASO-treated mice. Objective: 10x (upper) and 20x (lower), scale bars = 150 μm. Ordinary one-way ANOVA was used.

#### Supplementary Figure 4

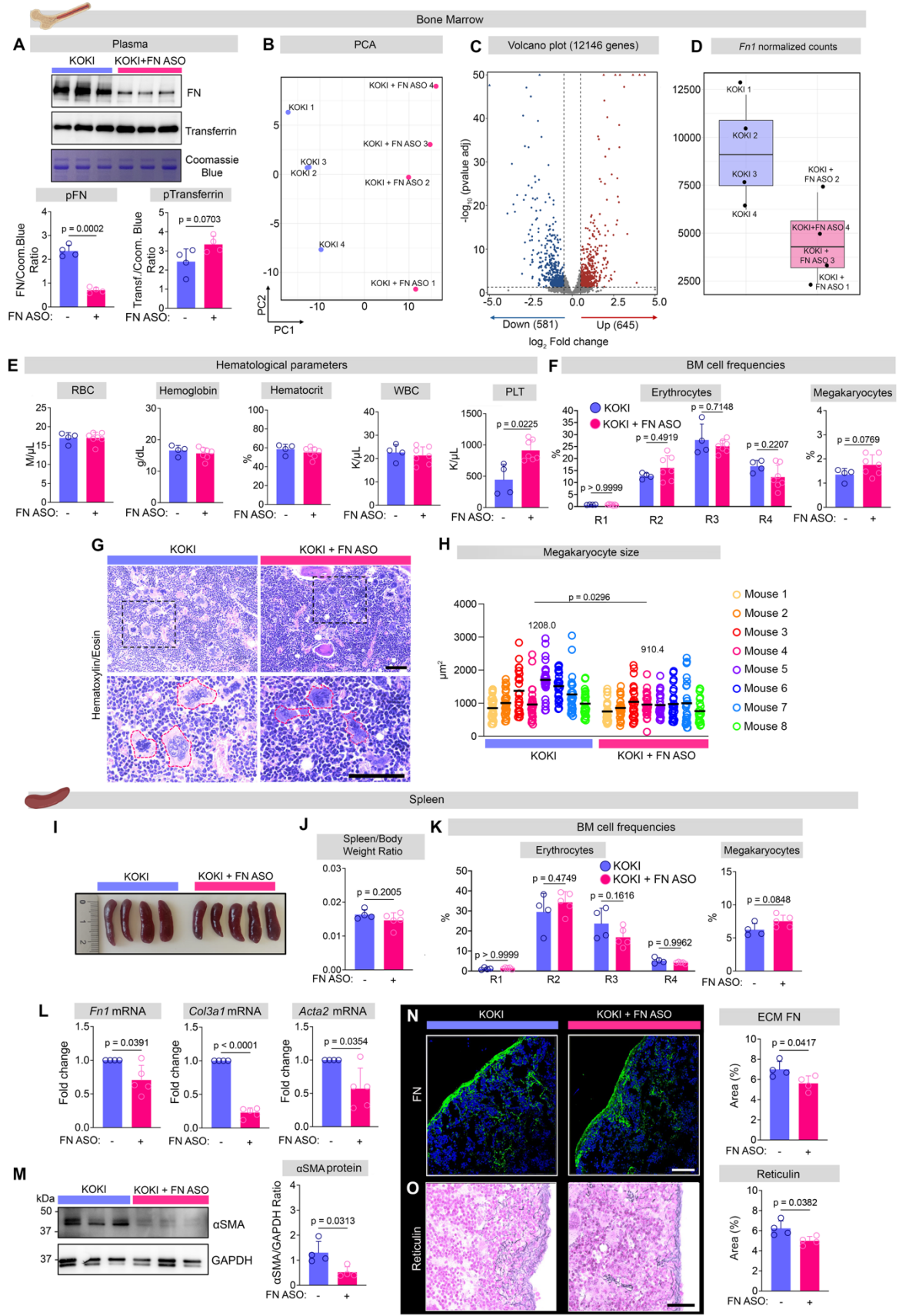

**Figure S4. FN ASO attenuates the pro-fibrotic phenotype of KOKI mice, with concomitant changes in the megakaryocytic/platelet lineage and reduction of FN expression and fibrosis-associated markers in spleens.**

**A)** Western blot analysis of plasma FN and transferrin and relative quantifications from KOKI+vehicle and KOKI+FN ASO mice (n=4). Coomassie Blue staining were used to ensure the equal loading. Welch's t test was used.

**B)** Principal Component Analysis (PCA) of BM samples from KOKI+vehicle and KOKI+FN ASO mice.

**C)** Volcano plot of differentially expressed genes of BM samples from KOKI+vehicle and KOKI+FN ASO mice.

**D)** Expression of *Fn1* in mouse RNA-seq data from vehicle- and FN ASO-treated KOKI mice (n = 4).

**E)** Hematological parameters of KOKI+vehicle and KOKI+FN ASO mice, including RBC count, hemoglobin, hematocrit, WBC count and PLTs (n = 4). Welch's t test was used.

**F)** Cell frequencies of erythrocytes and megakaryocytes from BM of KOKI+vehicle and KOKI+FN ASO (n = 4). R1= Proerythroblasts, R2= Basophilic erythroblasts, R3= Polychromatophilic erythroblasts, R4= Orthochromatophilic erythroblasts. Ordinary two-way ANOVA was used for erythrocytes; Welch's t test was applied to megakaryocytes.

**G-H)** Hematoxylin and eosin staining of BM sections from KOKI+vehicle and KOKI+FN ASO-treated mice (**G**) with analysis of megakaryocyte size (**H**). Objective: 10x (upper) and 30x (lower). Scale bars=250 μm. Each data point represents the mean value calculated from multiple cells/fields per mouse, with each mouse considered one biological replicate. Welch's t test was used.

**I-J)** Representative image of spleens from KOKI+vehicle and KOKI+FN ASO (**I**) and spleen weight normalized to body weight quantification (**J**). Welch's t test was used.

**K)** Cell frequencies of erythrocytes and megakaryocytes in spleens from KOKI+vehicle and KOKI+FN ASO. At least 4 replicates were analyzed. Ordinary two-way ANOVA was used for erythrocytes; Welch's t test was applied to megakaryocytes.

**L)** RT-qPCR analysis of *Fn1*, *Col3a1* and *Acta2* expression in spleens of KOKI+vehicle and KOKI+FN ASO. At least 4 replicates were evaluated. Welch's t test was used.

**M)** Western blot analysis of α-SMA expression in spleen cells from KOKI+vehicle and KOKI+FN ASO-treated mice, and its quantification. GAPDH was used as a loading control. At least 4 replicates were evaluated. Welch's t test was used.

**N)** Immunofluorescence analysis of FN in spleen sections from KOKI+vehicle and KOKI+FN

ASO-treated mice. Objective: 25x, scale bar = 150 μm. Welch's t test was used.

**O)** Reticulin staining in spleen sections from KOKI and KOKI+FN ASO-treated mice. Objective: 20x, scale bar = 100 μm. Welch's t test was used.

**Table S1:** Ranked list of differentially expressed genes in hMSCs treated with scrambled ASO, FN ASO, TGF-β1 plus scrambled ASO, or TGF-β1 plus FN ASO.

**Table S2:** Full ranked list of enriched pathways identified by pathway enrichment analysis in hMSCs treated with scrambled ASO, FN ASO, TGF-β1 plus scrambled ASO, or TGF-β1 plus FN ASO.

**Table S3:** Ranked list of differentially expressed genes in bone marrow cells from KOKI mice treated with vehicle or FN ASO.

**Table S4:** Full ranked list of enriched pathways identified by gene set enrichment analysis (GSEA) in bone marrow cells from KOKI mice treated with vehicle or FN ASO.
